# Targeting Pleiotrophin to mitigate high fat diet-induced liver metabolic disease: Insights into sex-specific metabolic protection

**DOI:** 10.64898/2025.12.02.691898

**Authors:** Héctor Cañeque-Rufo, Agata Zuccaro, Marta Inmaculada Sanz-Cuadrado, Federica Ruin, Ángela Olivera-Rodriguez, Javier Pizarro-Delgado, María Limones, Jimena Pita-Santibáñez, María Gracia Sánchez-Alonso, Julio Sevillano, Ángela M. Valverde, Esther Gramage, Gonzalo Herradón, María del Pilar Ramos-Álvarez

## Abstract

Obesity is a global health problem linked to the development of metabolic syndrome (MetS) and comorbidities such as metabolic dysfunction-associated steatotic liver disease (MASLD) and metabolic dysfunction-associated steatohepatitis (MASH). These diseases are characterised by systemic inflammation, lipid accumulation and tissue damage, which contribute to liver fibrosis and dysfunction. Pleiotrophin (PTN), a cytokine known for its role in tissue regeneration and energy metabolism, has emerged as a potential regulator of liver homeostasis. Herein, we demonstrate that *Ptn* deletion protects against body weight gain, metainflammation and high-fat diet (HFD)-induced MASLD and MASH development. Furthermore, our work uncovers the molecular mechanisms by which PTN may promote lipid synthesis and hepatic extracellular matrix remodelling. Results highlight PTN as a critical modulator of liver metabolism and systemic inflammation in the context of obesity, identifying it as a promising therapeutic target for the treatment of MASLD, MASH and related metabolic disorders, and point to a sexual dimorphism in adaptive metabolic strategies, with females demonstrating a greater degree of protection against the liver-damaging effects of diet-induced obesity.

**HIGHLIGHTS:**

- Females are more protected against the liver-damaging effects of the high-fat diet (HFD), suggesting a sexual dimorphism in adaptive metabolic strategies.
- PTN regulates lipid metabolism by promoting lipogenesis and lipid accumulation in liver cells through the activation of AKT and the inhibition of AMPKα; the absence of PTN reverses this process.
- Deletion of *Ptn* protects against high-fat diet (HFD)-induced weight gain, systemic inflammation, hepatic lipid accumulation, and the development of steatosis and liver fibrosis.
- PTN is a key modulator of metabolic, inflammatory and remodelling processes in the liver, and it is proposed as a promising therapeutic target for MASLD, MASH and other metabolic comorbidities.

**Graphical abstract.**
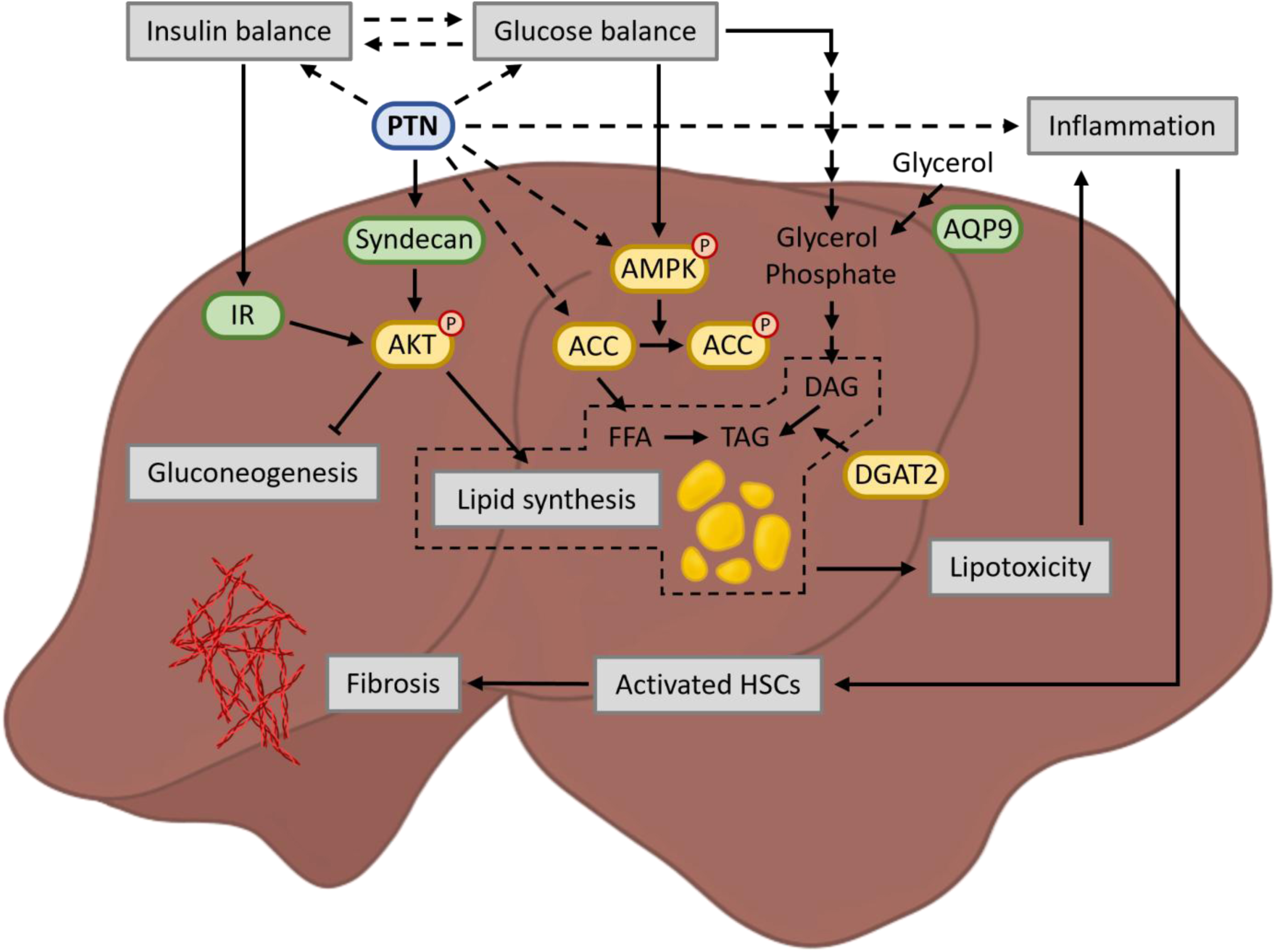
PTN liver signalling pathway. acetyl-CoA carboxylase (ACC); 5’ adenosine monophosphate-activated protein kinase (AMPK); protein kinase B (AKT); aquaporin 9 (AQP9); diacylglycerides (DAG); diacylglycerol O-acyltransferase 2 (DGAT2); free fatty acids (FFA); hepatic stellate cell (HSC); insulin receptor (IR); pleiotrophin (PTN); triacylglycerides (TAG).

## INTRODUCTION

According to the latest World Health Organisation report, obesity has become one of the most prevalent diseases worldwide [1]. This chronic disease is associated with the development of metabolic syndrome (MetS) [2], a complex disorder characterised by a set of cardiovascular and metabolic alterations, including hypertension, dyslipidaemia, fasting hyperglycemia, and hyperinsulinemia, among others [3]. All these alterations lead to a state of systemic inflammation, known as metainflammation, which contributes to the development of diseases such as type 2 diabetes mellitus, metabolic dysfunction-associated steatotic liver disease (MASLD) and even neurodegenerative diseases such as Alzheimer’s disease [2, 4–8]. Among the comorbidities associated with obesity, MASLD is one of the most detrimental. Excessive lipid accumulation within hepatocytes results in hepatic steatosis [9]. This condition can present as MASLD, which includes benign forms of the disease such as simple fatty infiltration. However, when fat accumulation triggers lipotoxicity, accompanied by immune cell infiltration into the liver parenchyma and the release of pro-inflammatory cytokines responsible for cell damage, the condition progresses to metabolic dysfunction-associated steatohepatitis (MASH) [9–11]. In this context, the presence of MetS significantly increases the risk of simple steatosis progressing to MASH, where the liver ceases to function properly [9, 12]. Notably, several clinical studies have reported that premenopausal women exhibit a lower prevalence and severity of liver fibrosis compared to men, suggesting a protective role of estrogens that diminishes after menopause [13, 14]. Although multiple pathways have been proposed, the precise molecular mechanisms underlying this sex-based protection against liver fibrosis remain incompletely understood. The resulting liver damage leads to alterations in peripheral metabolism that ultimately affect other organs [12]. All the comorbidities associated with obesity lead to a collapse of health systems and large investments in their treatment. Therefore, there is an urgent need to investigate therapeutic targets for this disease and its associated alterations.

Pleiotrophin (PTN) is a cytokine mainly expressed in the central nervous system (CNS) [15]. Furthermore, its expression is strictly limited in adulthood, except for processes associated with tissue regeneration, cell growth and angiogenesis [16–19]. Previous studies from our group have pointed out the pivotal role of PTN in regulating insulin sensitivity, energy metabolism and thermogenesis [20, 21]. In the liver, PTN is expressed in the adult stage and plays a mitogenic role in the development and regeneration of liver tissue [22, 23]. However, its function in hepatic metabolism and its involvement in the progression of MASLD have not been fully elucidated. Our recent findings suggest that *Ptn* deletion may confer protection against hepatic steatosis, as female mice subjected to a 45% high-fat diet (HFD) for three months exhibited reduced lipid accumulation compared to their wild-type counterparts [21]. However, several questions remain open, including whether similar protective effects occur in male mice and whether *Ptn* deletion also mitigates other liver disturbances commonly associated with steatosis. Therefore, the present study aimed to investigate the involvement of PTN in hepatic metabolism, under both physiological and pathological conditions, using HFD-induced obesity and *Ptn* genetically deficient (*Ptn*^−/−^) mice of both sexes. In addition, *in vitro* studies in liver cell lines were performed to determine the molecular mechanisms involved in the observations *in vivo*. In this context, our study highlights the pivotal role of endogenous PTN in body weight gain and metabolic changes associated with diet-induced obesity. Our results demonstrate that genetic deletion of *Ptn* significantly prevents the development of metainflammation, hepatic lipid accumulation and subsequent HFD-induced liver pathologies such as steatosis and fibrosis. Importantly, this study highlights significant sex-dependent differences in the progression of HFD-induced liver alterations, with females exhibiting hepatic protection compared to males. Understanding the molecular basis of these sex-specific responses is crucial for developing targeted therapies for metabolic liver diseases. In addition, our data reveal the role of PTN in liver metabolism under physiological and pathological conditions, emphasising its importance for metabolic homeostasis. Results also position PTN as a promising therapeutic target for the treatment of obesity and its comorbidities, opening new avenues for clinical intervention in metabolic diseases.

## MATERIALS AND METHODS

### In vivo studies

#### Animals and samples collection

*Ptn*^−/−^ mice were generated as previously described [24]. Male and female C57BL/6J wild-type (*Ptn*^+/+^) and *Ptn*^−/−^ mice were randomly divided and housed in a specific pathogen-free room at 22 ± 1°C with 12h light/dark cycles, with free access to water and randomly fed *ad libitum* with standard chow diet (STD, 18 kcal% fat, 58 kcal% carbohydrates, and 24% kcal protein; 3.1 kcal·g^−1^) or a HFD (D12492, 60 kcal% fat, 20 kcal% carbohydrates, and 20% kcal protein; 5.24 kcal·g^−1^) as corresponding. The diet intervention started at 3 months of age and was maintained for 6 months. All the animals were handled and maintained in accordance with the European Union Laboratory Animal Care Rules (2010/63/EU directive), and protocols were approved by the Animal Research Committee of CEU San Pablo University and the Comunidad de Madrid (PROEX 140.3/22).

After 6h of fasting, 9-month-old mice from each experimental group (n = 7/group) were sacrificed by decapitation. The livers were removed and post-fixed in paraformaldehyde (PFA) 4% for 48h at 4°C for histological analysis. Fixed livers were dehydrated and embedded in paraffin and sectioned at 3 μm thickness with a microtome (Leica Biosystems). For molecular studies, the livers were collected and stored at -80 °C. Blood samples were collected in EDTA-containing tubes, and after centrifugation at 1500 rpm for 10 minutes, the plasma was obtained and stored at -20°C until analysis.

#### Pyruvate tolerance test

To perform the pyruvate tolerance test (PTT), mice from each condition (treated with the corresponding diet for 3 months) were fasted for 16h and then injected intraperitoneally with sodium pyruvate (2 g/kg body weight) dissolved in saline. Blood glucose levels were measured from the tail vein at the indicated times using a glucometer (Contour, Bayer, Basel, Switzerland).

#### Histology and image analysis

Two randomly selected liver sections per animal were deparaffinised, rehydrated and stained with haematoxylin and eosin (H&E) to examine the tissue morphology and lipid accumulation. Additional sections were stained with picrosirius red to examine collagen and fibrosis. The sections were analysed by optical microscopy (Leica Biosystems, Barcelona, Spain). Analysis was performed using *Fiji* software (NIH, Bethesda, MD, USA, Version 1.50f). For H&E staining, five images (10X objective) from each mouse were taken to estimate the degree of hepatic steatosis by examining the percentage of hepatocytes with lipid accumulation and ballooned hepatocytes. In the experimental groups where a clear state of steatosis was observed, the pattern of lipid accumulation was studied. A distinction was made between macrovesicular and microvesicular accumulation. Additionally, to study the degree of fibrosis, at least 10 blood vessels from each experimental condition were analyzed in bright field for picrosirius red staining (10X objective) to determine the collagen surrounding them. Subsequently, type I and type III collagen were determined using polarised light.

#### Hepatic lipid analysis

Total lipids were extracted following the Folch method [25]. Then, the different lipid fractions were separated by thin-layer chromatography in Silicagel for quantification as previously described [26].

#### Circulating proteins and inflammatory cytokines analysis

Total plasma protein concentration was measured using the Pierce BCA protein method (Thermo-Scientific, Waltham, MA, USA), and the Helena, SAS-MX SP-10 kit was used to detect changes in the profile of the major circulating proteins. For that, after protein separation by agarose gel electrophoresis in Tris-barbital buffer, the gels were stained with Paragon blue 0.5% in 5% acetic acid solution and then scanned for quantification.

Plasma inflammatory markers were assessed using the commercially available Olink® Target 96 panels (Uppsala, Sweden). In brief, the target protein binds with high specificity to the double oligonucleotide-labelled antibody probe. Then, microfluidic real-time PCR amplification of the oligonucleotide sequence is used to quantitatively detect the resulting DNA sequence. Using internal and external controls, the resulting threshold cycle (Ct)-data are processed for quality control and normalised. Normalised protein expression (NPX) values, an arbitrary unit of log2 scale corresponding to higher protein levels, were expressed as fold change. One control sample that failed to pass the quality control, was excluded from further analysis.

#### Tissue protein extraction and immunoblotting

Fifty mg of liver tissue was homogenized in lysis buffer (pH = 7.4) containing 30 mM HEPES, 5 mM EDTA, 1% Triton X-100, 0.5% sodium deoxycholate, 8 mM Na3VO4, 1 mM NaF, and 2 mM of the protease inhibitor mixture Pefabloc (Roche Diagnostics, Barcelona, Spain). Protein concentration was measured using the Pierce BCA protein method (Thermo-Scientific, Waltham, MA, USA). An equal amount of protein was subjected to SDS-PAGE electrophoresis, transferred to PVDF membranes (Amersham-GE Healthcare, Amersham, UK), and incubated with anti-aquaporin 9 (AQP9) (Invitrogen; #PA5-76416; 1:500; rabbit), anti-diacylglycerol O-acyltransferase 2 (DGAT2) (Thermo Fisher; #PA5-103785; 1:1000; rabbit), anti-AMP-activated protein kinase α (AMPKα) (Cell Signaling; #2532; 1:500; rabbit), anti-phospho-AMPKα (Thr172) (Cell Signaling; #2531; 1:1000; rabbit), anti-acetyl-CoA carboxylase (ACC) (Millipore; #MABS2024; 1:500; mouse), and anti-phospho-ACC (pACC) (Ser79) (Millipore; #07-303; 1:500; rabbit) antibodies. Vinculin (Sigma-Aldrich; #V9264; 1:10000; mouse) was used as a loading control. After the incubation with the corresponding horseradish peroxidase-conjugated secondary antibody, proteins were visualized by the enhanced chemiluminescence (ECL) system (Amersham; #RPN2236; UK) using the ChemiDoc Imaging system (Bio-Rad, Hercules, CA, USA). Levels of proteins were quantified by densitometry in each animal sample using Image Lab image acquisition and analysis software (Bio-Rad, Hercules, CA).

#### Tissue quantitative real-time PCR

Total RNA from liver was isolated using the Total RNA Isolation Kit (Nzytech, Lisbon, Portugal). First-strand cDNA was synthesized using the first-strand cDNA Synthesis Kit (Nzytech, Lisbon, Portugal), and 5 μg of RNA were reverse-transcribed to DNA. Quantitative real-time PCR (qPCR) analysis was performed using the SYBR green method (Quantimix Easy kit, Biotools, Madrid, Spain) in a CFX96 Real-Time System (Bio-Rad, Hercules, CA, USA). The relative expression of each gene was normalized using actin (*Actb*) and ribosomal protein L13 (*Rpl13*) as housekeeping genes, and the data were analyzed by the Livak method. The primer sequences used are shown in **Table 1**.

**Table 1.**
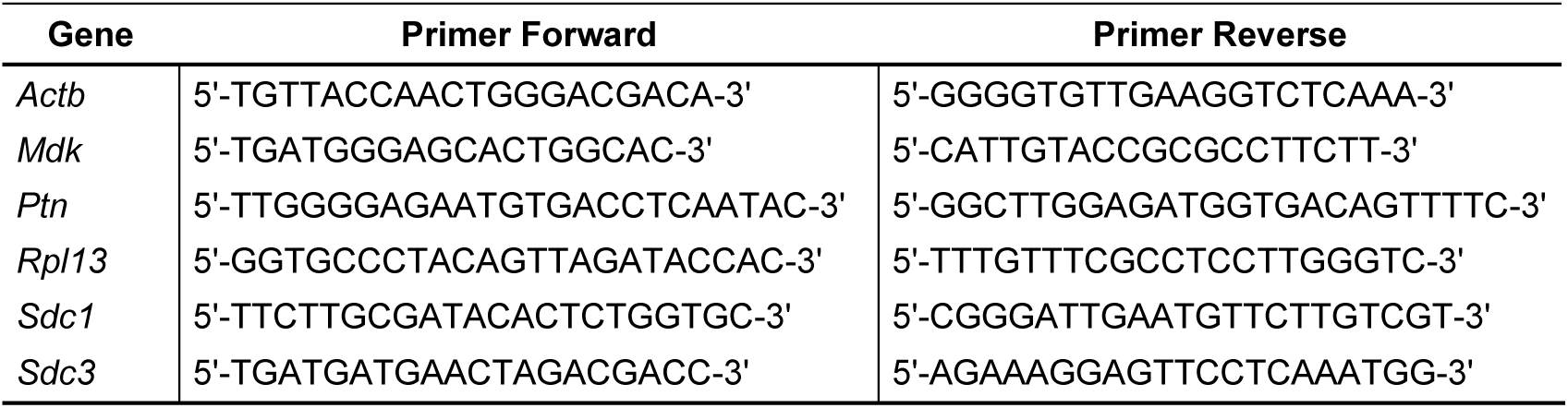
Primer sets are used for tissue qPCR analysis. *Actb*: Actin beta; *Mdk*: Midkine; *Ptn*: Pleiotrophin; *Rpl13*: Ribosomal protein L13; *Sdc1*: Syndecan 1; *Sdc3*: Syndecan 3.

### In vitro studies

#### Cell culture and treatments

The hepatocellular carcinoma Huh-7 cell line was cultured at 37°C and 5% CO2 and grown in DMEM (Dulbecco’s Modified Eagle Medium) High Glucose (4,5 g/L) with L-glutamine (Biowest), supplemented with 10% (vol./vol.) fetal bovine serum (FBS) and 1% (vol./vol.) penicillin/streptomycin (Sigma-Aldrich). For the different treatments, the cells were starved by decreasing the FBS to 3%. The effect of PTN was tested by incubating the cells with 0,5 µg/mL of recombinant PTN (R&D Systems) for 0, 0.25, 0.5, 1, 2 or 4 h. For analysis of the cellular lipid content, cells were treated with 10 nM human recombinant insulin, 0.5 µg/mL of recombinant PTN, or a combination of both for 48 h.

#### Cell protein extraction and immunoblotting

Protein extraction, concentration, electrophoresis and transfer were performed as described above. Membranes were incubated with anti-protein kinase B (AKT) (Cell Signaling; #CS9272; 1:1000; rabbit), anti-phospho-protein kinase B (pAKT) (Ser473) (Cell Signaling; #CS4060; 1:2000; rabbit), anti-ACC (Sigma-Aldrich; #05-1098; 1:500; mouse), and anti-pACC (Ser79) (Cell Signaling; #CS3661; 1:500; rabbit) antibodies.

#### Intracellular lipid analysis

Oil Red O (OR) stain was used to evaluate the intracellular lipid content. A stock solution of OR was prepared by dissolving 0.3 g of OR powder (Sigma-Aldrich) in 100 mL of 2-propanol. The stock solution was filtered and stored at room temperature. The working solution was prepared freshly for each experiment by diluting 3 parts of the stock solution with 2 parts of distilled water, and it was filtered immediately before use. According to the staining protocol, cells were washed twice with PBS 1X and fixed with 4% PFA for 30 min after the treatment. The cells were washed twice with distilled water and incubated with 60% 2-propanol for 5 min. After discarding the isopropanol, cell monolayers were stained with OR working solution for 15 min at room temperature. Cells were analysed by optical microscopy (Leica Biosystems, Barcelona, Spain). Analysis was performed using *Fiji* software (NIH, Bethesda, MD, USA, version 1.50f). Images (20X objective) from each well were acquired to estimate the area stained with OR and normalised by cell number.

For the analysis of triacylglycerides (TAG) cell content, at the end of the treatment period, the cell monolayers were washed twice with cold PBS 1X. Cells were then homogenised in 100 µL of 5% NP-40/ddH2O solution. The samples were heated at 95°C for 5 min until the NP-40 solution became cloudy and then cooled down to room temperature. This step was repeated twice to solubilise all TAG. The samples were then frozen at -80°C for 6 hours. TAG were quantified by the glycerol-3-phosphate/peroxidase chromogenic method using a Spinreact kit, according to the manufactureŕs instructions in 5 µL of each sample; after incubation for 5 min at 37°C with 250 µL of the working reactive, the absorbance was measured at 505 nm on a SPECTROstarNano microplate reader (BMG Labtech, Germany).

#### Cell quantitative real-time PCR

Total RNA isolation, first-strand cDNA synthesis, and qPCR analysis were performed as described above. The relative expression of each gene was normalized using glyceraldehyde-3-phosphate dehydrogenase (*GAPDH*) and ribosomal protein L30 (*RPL30*) as housekeeping genes, and the data were analyzed by the Livak method. The primer sequences used are shown in **Table 2**.

**Table 2.**
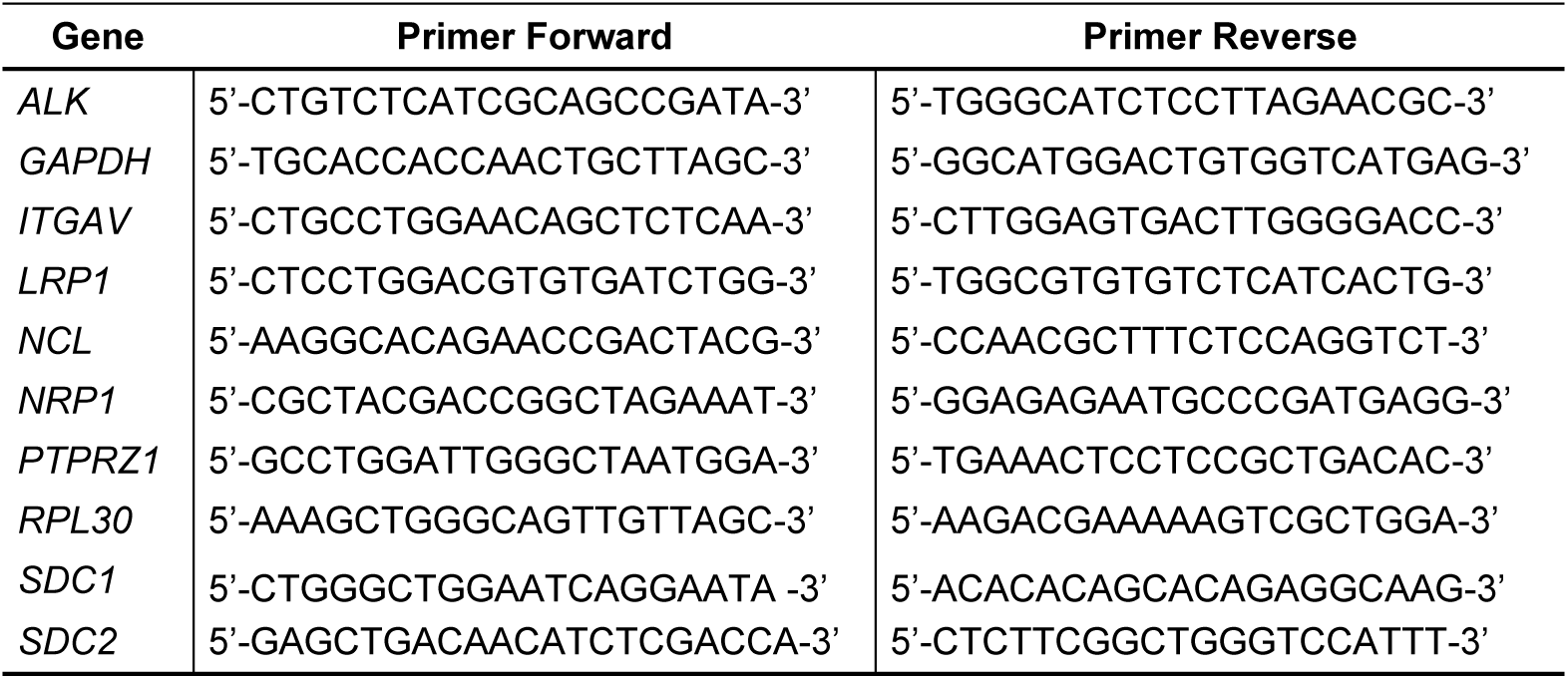
Primer sets are used for cell qPCR analysis. *ALK*: ALK receptor tyrosine kinase; *GAPDH*: glyceraldehyde-3-phosphate dehydrogenase; *ITGAV*: integrin subunit alpha V; *LRP1*: LDL receptor related protein 1; *NCL*: nucleolin; *NRP1*: Neuropilin 1; *PTPRZ1*: Protein tyrosine phosphatase receptor type Z1; *RLP30*: ribosomal protein L30; *SDC1*: Syndecan 1; *SDC2*: Syndecan 2.

#### Statistical Analysis

Statistical analyses were performed using IBM-SPSS v.25 software. The mean value and the standard error of the mean (±SEM) are presented in the graphs. The Shapiro-Wilk test was used to test the normality of the sample distribution. If the normal distribution of the data was confirmed, parametric tests were used. Specifically, the increment of body weight over time and food intake were analysed using a three-way repeated measures ANOVA test with sex, genotype, and diet as variables. In the analysis of vesicularity, as only two experimental groups were compared, a T-Student test was used. For the evaluation of the response to insulin or PTN in the cell line, two- or one-way ANOVA test was applied as correspond. For the remaining studies, data were analysed using a three-way ANOVA test with the same variables. When relevant, a two-way ANOVA we used to better dissect the effect of each variable, excluding the non-significant variable when the three-way ANOVA results allowed. When normal distribution of the data was not confirmed, non-parametric tests were used. Specifically, the pre-albumin fraction and the percentage of ballooned hepatocytes were analysed using a Kruskal-Wallis test with sex, genotype, and diet as variables. Bonferroni’s *post-hoc* analysis was used to compare the differences between groups when an interaction between variables was found. All references to statistical significance are made to the three-way or two-way ANOVA test’s individual factors and their interaction or to the Kruskal-Wallis test. In the figures corresponding to the *in vivo* studies, significant differences are represented with (*) for differences between diets, (#) for differences between genotypes, and (&) for differences between sexes. For the figures corresponding to the *in vitro* studies, significant differences are represented with (*) for the treatment effects and (#) for differences between treatments. A 95% confidence interval was used for all statistical comparisons. All results of the statistical analysis of the ANOVAs are shown as supplementary material.

## RESULTS

### *Ptn* deletion protects against high-fat diet induced obesity

First, the body weight of the animals was monitored throughout the experiment. Two-way repeated measures ANOVA revealed that both genotype and diet significantly affected the increment of body weight over time in both males (**Figure 1A**) and females (**Figure 1B**). In addition, a significant interaction between variables was found in both sexes. Moreover, three-way ANOVA revealed that genotype, diet, and sex, significantly influenced the final body weight, although we did not detect significant interactions between variables (**Figure 1C**). We then examined the cumulative food intake throughout the experiment to discern whether the lack of weight gain was related to lower intake. Two-way repeated measures ANOVA showed that diet significantly decreased the cumulative food intake in male and a tendency to decrease in female mice (**Supplementary Figure 1A-B**). No effects of genotype on daily calorie intake (**Figure 1D**) or in cumulative food consumption were observed (**Supplementary Figure 1**). Although male *Ptn*^−/−^ mice initially showed increased body weight when fed an HFD, this did not affect the final gain. However, this effect was not observed in female *Ptn*^−/−^ animals. The results indicate that the body weight gain caused by the HFD was significantly lower in *Ptn*^−/−^ than in *Ptn*^+/+^ animals (statistics in **Supplementary Table 1A and I**).

**Figure 1.**
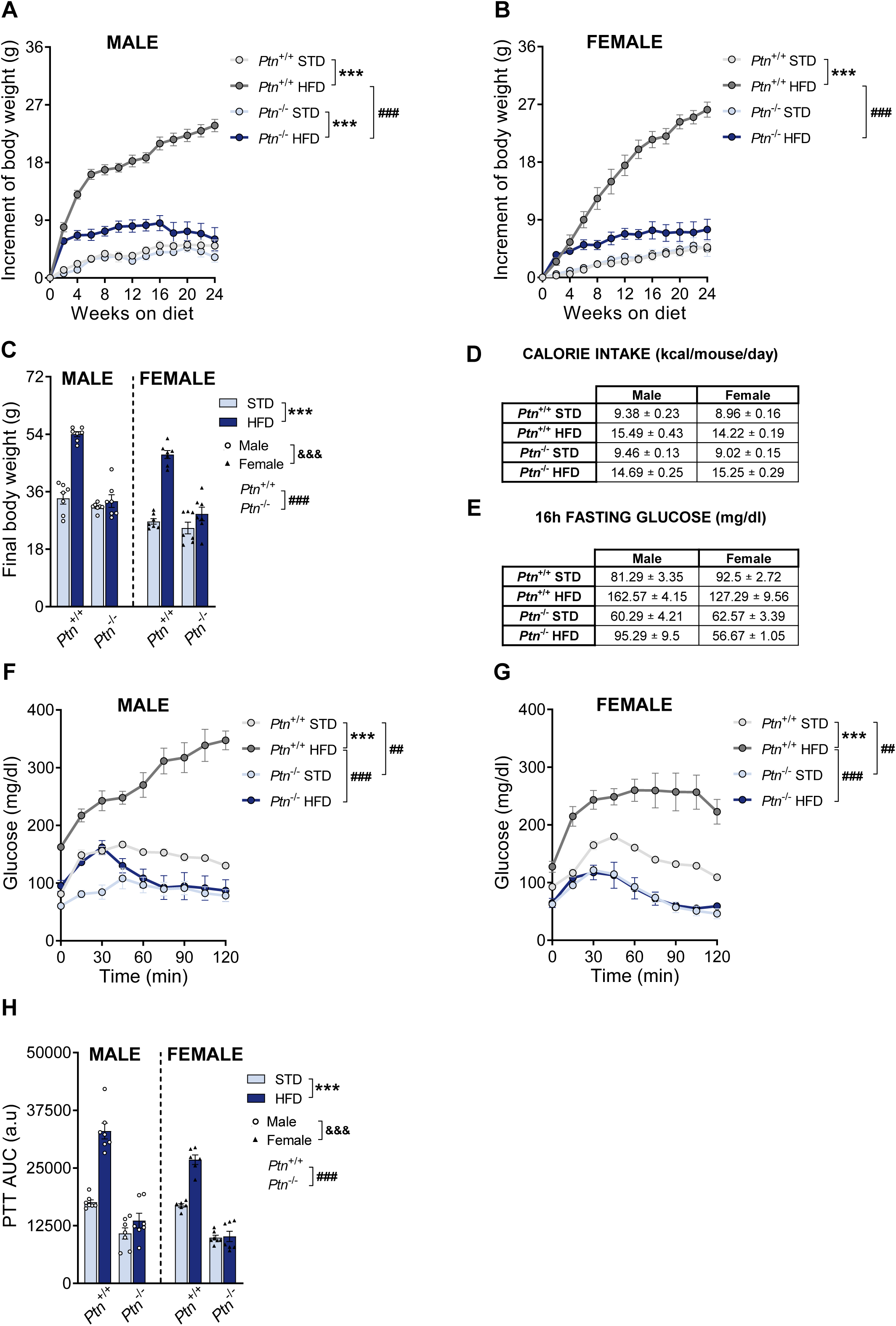
High-fat diet feeding and *Ptn* deletion effects on body weight and gluconeogenesis from piruvate. Increment of body weight over time in male (A) and female (B), final body weight (C) and daily calorie intake (D) in *Ptn*^+/+^ and *Ptn*^−/−^ mice fed with a STD or HFD for 6 months. Basal glucose after 16h fasting (E); glycemia during the pyruvate tolerance test (PTT) in male (F) and female (G), and area under the curve from the PTT (H) in *Ptn*^+/+^ and *Ptn*^−/−^ mice fed with a STD or HFD for 3 months. Data are presented as mean ± SEM (n = 7 animals per group). Differences between diets are represented with: \*\*\**p* < 0.001; between genotypes with: ^##^*p* < 0.01; ^###^*p* < 0.001; and between sexes with: ^&&&^*p* < 0.001.

### High-fat diet and *Ptn* deletion impair basal glycemia and gluconeogenesis

To study hepatic gluconeogenesis, we performed a pyruvate tolerance test (PTT). Due to the prolonged fasting required for this test and the severe damage produced by the HFD after six months, there were numerous casualties in the experimental groups of *Ptn*^+/+^-HFD males and females. For this reason, the test was only carried out after three months with the HFD. First, basal glycemia after 16h of fasting was determined (**Figure 1E**). Three-way ANOVA revealed that genotype, diet and sex significantly altered the basal circulating glucose. However, no interaction between the three variables was found. HFD increased fasting glycemia in *Ptn*^+/+^ animals, more markedly in males than in females. In addition, *Ptn*^−/−^ mice had lower basal glucose under STD conditions. Strikingly, HFD produced a smaller increase in basal glucose levels in male *Ptn*^−/−^ compared with *Ptn*^+/+^ mice, whereas female *Ptn*^−/−^ mice fed HFD were fully protected against these alterations.

Next, PTT was performed. Two-way repeated measures ANOVA revealed that both genotype and diet significantly affected the blood glucose after pyruvate injection in both males (**Figure 1F**) and females (**Figure 1G**). In addition, a significant interaction between variables was found in both sexes. HFD diet results in a significant rise in glycemia following pyruvate injection, and whereas in females this effect tends to normalise over time, this is not the case in males. Interestingly, *Ptn*^−/−^ animals showed lower blood glucose after pyruvate injection in all cases, and this was unaffected by the HFD diet, being even, in females, the same in those fed the HFD as in those fed the STD diet.

After studying the changes over time in blood glucose after pyruvate injection, the area under the curve (AUC) was calculated in order to study the magnitude of the observed effect. Three-way ANOVA indicated that genotype, diet and sex significantly altered the AUC from PTT, although no interaction between the three variables was found (**Figure 1H**) (statistics in **Supplementary Table 1A**).

### *Ptn*, *Mdk* and *Sdc* mRNA in the liver are differentially regulated by sex, genotype and diet

Next, we examined the mRNA of *Ptn*, midkine (*Mdk*), syndecan 1 (*Sdc1*) and syndecan 3 (*Sdc3*) in the liver. Two-way ANOVA showed that neither diet nor sex significantly affected the amount of *Ptn* mRNA. However, a significant interaction between variables was found (**Figure 2A**). *Post-hoc* analysis indicated that HFD increased *Ptn* mRNA levels only in male *Ptn*^+/+^ mice. Three-way ANOVA showed a significant effect of genotype and sex, but not diet, on *Mdk* mRNA. However, a significant interaction between three variables was found (**Figure 2B**). Overall, HFD increased the *Mdk* mRNA expression in male *Ptn*^+/+^ but not in male *Ptn*^−/−^ mice. Interestingly, those changes were not found in female mice that showed higher *Mdk* mRNA compared with male mice. Finally, three-way ANOVA revealed that genotype, diet, and sex significantly affected the *Sdc1* mRNA (**Figure 2C**), whereas diet and sex, but not genotype, significantly modified the *Sdc3* mRNA expression (**Figure 2D**). In addition, no interaction between the three variables was found for either *Sdc1* or *Sdc3* (statistics in **Supplementary Table 1B**).

**Figure 2.**
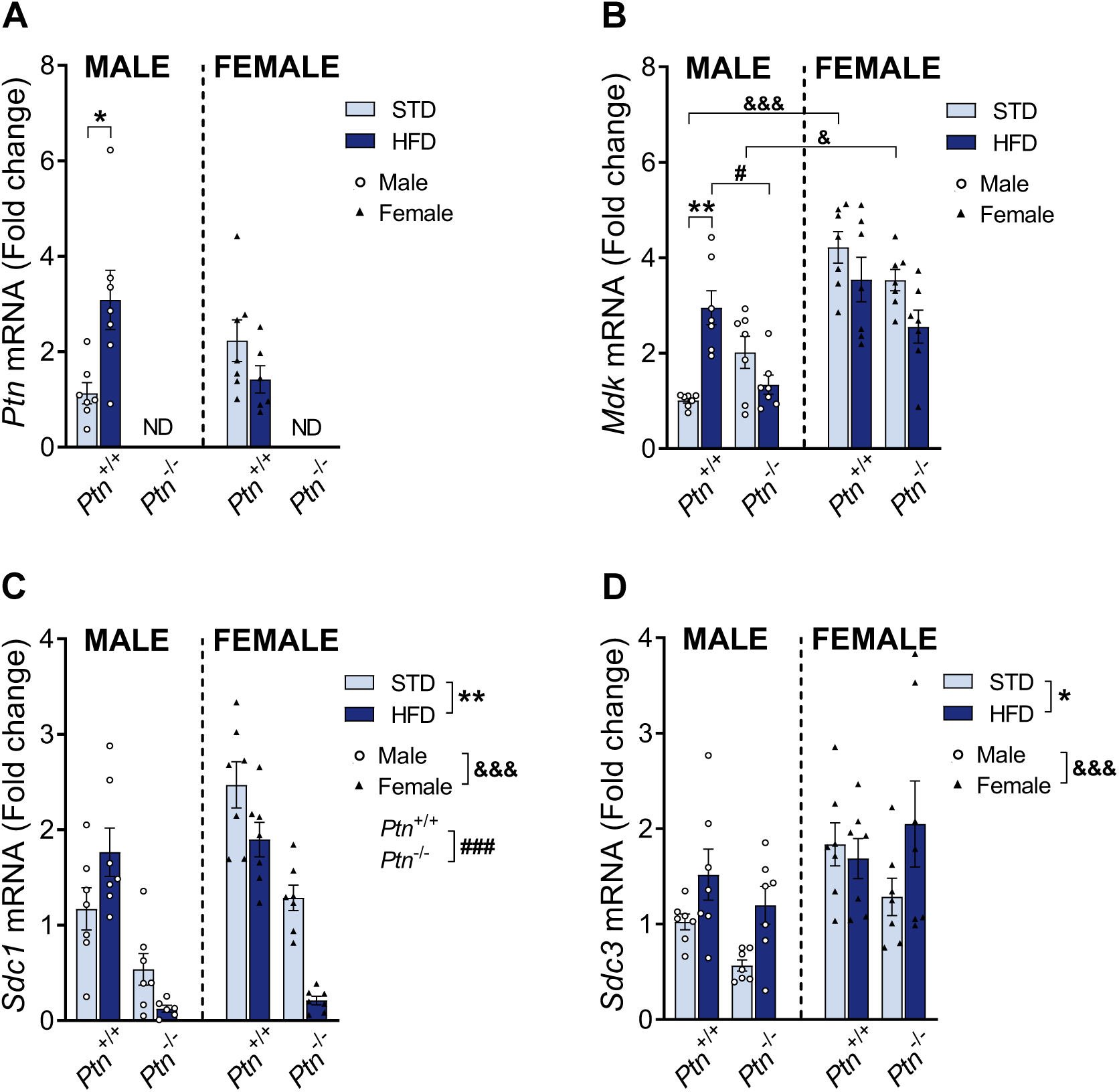
Hepatic *mRNA* of *Ptn, Mdk, Sdc1* and *Sdc3* is modulated by genotype, diet, and sex. Pleiotrophin (*Ptn*) mRNA (A); midkine (*Mdk*) mRNA (B); syndecan 1 (*Sdc1*) mRNA (C); syndecan 3 (*Sdc3*) mRNA (D) in the liver of male and female *Ptn*^+/+^ and *Ptn*^−/−^ mice fed with a STD or HFD for 6 months. Data are presented as mean ± SEM (n = 7 animals per group; circles correspond to males and triangles correspond to females). Differences between diets are represented with: \**p* < 0.05; \*\**p* < 0.01; between genotypes with: #p < 0.05; ###p < 0.001; and between sexes with: ^&^*p* < 0.05; ^&&&^*p* < 0.001.

### *Ptn* deficiency protects against high-fat diet induced systemic inflammation

Total plasma proteins and major protein fractions were analysed as liver markers. Three-way ANOVA revealed that HFD significantly increased the total circulating proteins. However, a significant interaction between three variables was found (**Figure 3A**). Specifically, we found that HFD increased the protein concentration in male *Ptn*^+/+^ but not in *Ptn*^−/−^ mice. Then, we studied the relative percentage of the protein fractions in each group (**Figure 3B**). Kruskal-Wallis test indicated that genotype and diet, but not sex, significantly affected the pre-albumin relative percentage. Next, three-way ANOVA showed that genotype significantly affected the α1-globulin relative percentage; diet significantly affected the albumin, α1-globulin, and α2-globulin relative percentages; and sex significantly affected the α1-globulin, α2-globulin, and γ-globulin relative percentages. In addition, a significant interaction between three variables was found in the case of α2-globulin relative percentage. In brief, HFD increased α1-globulin and α2-globulin and decreased the albumin relative percentages only in *Ptn*^+/+^ animals. In addition, we observed that the γ-globulin relative percentage was increased in female mice.

**Figure 3.**
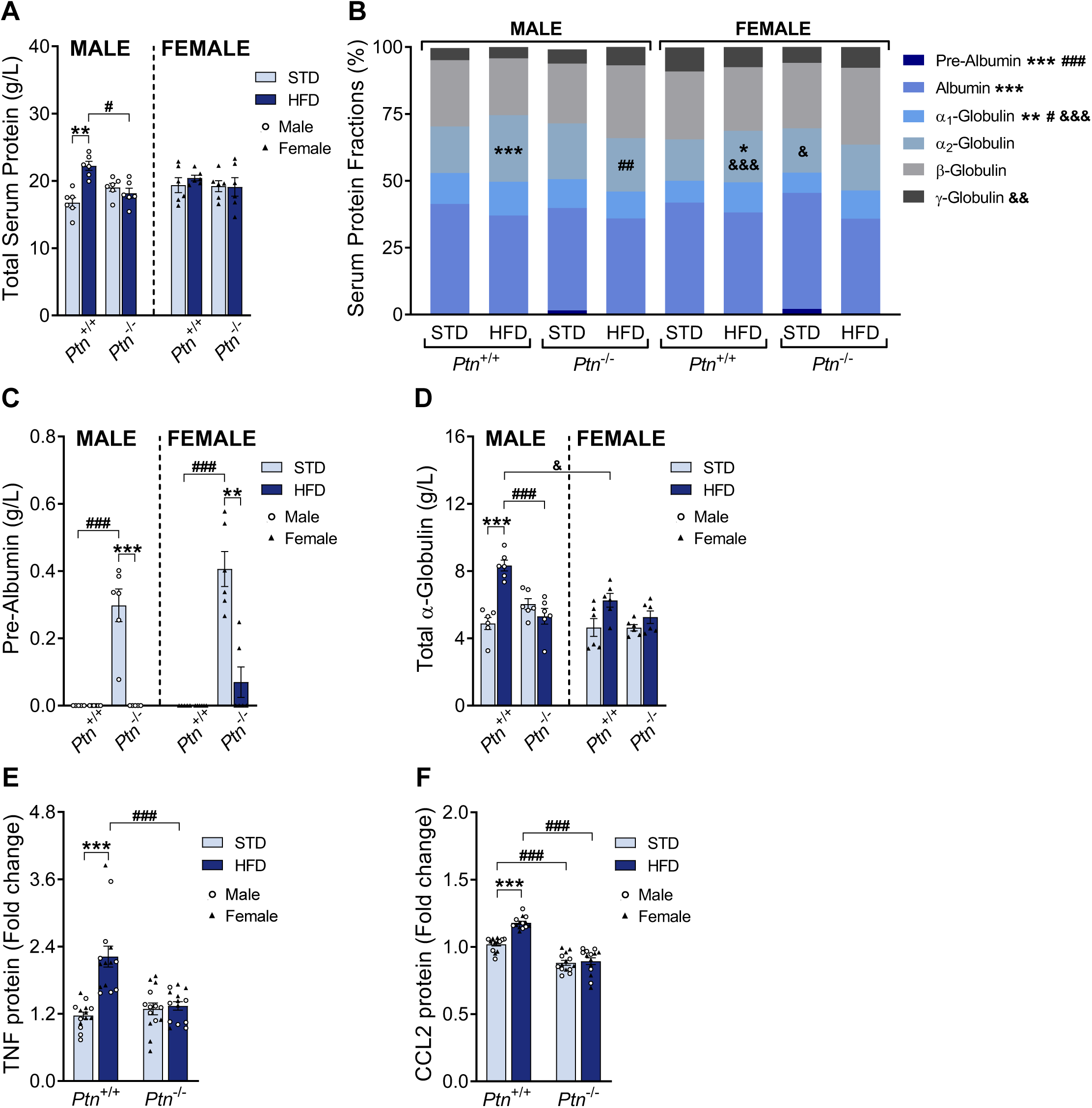
High-fat diet feeding and *Ptn* deletion effects on serum proteins and circulating proinflammatory molecules. Total serum proteins (A) and % of serum protein fractions (B). Serum pre-albumin (C); total α-globulin proteins (D); tumor necrosis factor (TNF) (E) and C-C motif chemokine 2 (CCL2) (F) in male and female *Ptn*^+/+^ and *Ptn*^−/−^ mice fed with a STD or HFD for 6 months. Data are presented as mean ± SEM (n = 6-7 animals per group; circles correspond to males and triangles correspond to females). Differences between diets are represented with: \**p* < 0.05; \*\**p* < 0.01; \*\*\**p* < 0.001; between genotypes with: ^#^*p* < 0.05; ^##^*p* < 0.01; ^###^*p* < 0.001; and between sexes with: ^&^*p* < 0.05; ^&&^*p* < 0.01; ^&&&^*p* < 0.001.

We observed similar results when the concentration of each protein fraction was calculated. Kruskal-Wallis test indicated that genotype and diet, but not sex, significantly affected the circulating concentration of pre-albumin (**Figure 3C**). In addition, three-way ANOVA revealed that genotype significantly affected the α1-globulin, α2-globulin; diet significantly affected the α1-globulin and α2-globulin; and sex significantly affected the α1-globulin, α2-globulin, and γ-globulin. Moreover, a significant interaction between three variables was found in the case of α2-globulin (**Supplementary Figure 2**). Finally, three-way ANOVA showed that genotype, diet, and sex significantly affected the concentration of the total α-globulin fraction and a significant interaction between variables was found (**Figure 3D**). In brief, we observed that *Ptn*^+/+^ animals fed a HFD showed an increase in α-globulin, an effect that does not occur in *Ptn*^−/−^ animals fed HFD. In addition, we found that γ-globulin was increased in female *Ptn*^+/+^ mice (**Supplementary Figure 2E**). Finally, the presence of pre-albumin was particularly striking, as it was observed only in *Ptn*^−/−^ animals and decreased with HFD (**Figure 3C**).

To complete the analysis of the effects of HFD and PTN deletion on metainflammation, proinflammatory molecules were analysed in plasma. Three-way ANOVA revealed that genotype and diet, but not sex, significantly affected the circulating concentration of tumour necrosis factor (TNF) (**Figure 3E**) and C-C motif chemokine 2 (CCL2) (**Figure 3F**). In addition, no interaction between the three variables was found. To further explore the effect of genotype and diet, we employed two-way ANOVA excluding the sex variable. Again, we found a significant effect of genotype and diet in both circulating TNF and CCL2. A significant interaction between genotype and diet was found in both cases. These results indicate that HFD increases plasma levels of TNF and CCL2 in *Ptn*^+/+^ mice, but not in *Ptn*^−/−^ animals. Furthermore, this animal model showed lower plasma levels of CCL2 at baseline (statistics in **Supplementary Table 1C and J**).

### *Ptn* deficiency prevents the development of hepatic steatosis induced by high-fat diet

Next, we studied the effects of HFD and *Ptn* deletion on the liver. Three-way ANOVA revealed that genotype, diet, and sex significantly affected the liver weight, and a significant interaction between the three variables was found (**Figure 4A**). In addition, three-way ANOVA showed that genotype, diet, and sex significantly modified the relative liver weight, and a significant interaction between the three variables was found (**Figure 4B**). Overall, we found that HFD produces an increase in liver weight only in *Ptn*^+/+^ mice, which was more pronounced in male than in female *Ptn*^+/+^ animals. Furthermore, when normalising liver weight to body weight, we found that the percentage was increased only in male *Ptn*^+/+^ animals after HFD intake. In addition, *Ptn*^−/−^ animals, regardless of diet, had a lower relative liver weight than *Ptn*^+/+^ animals, suggesting that *Ptn* deletion is associated with reduced liver growth.

**Figure 4.**
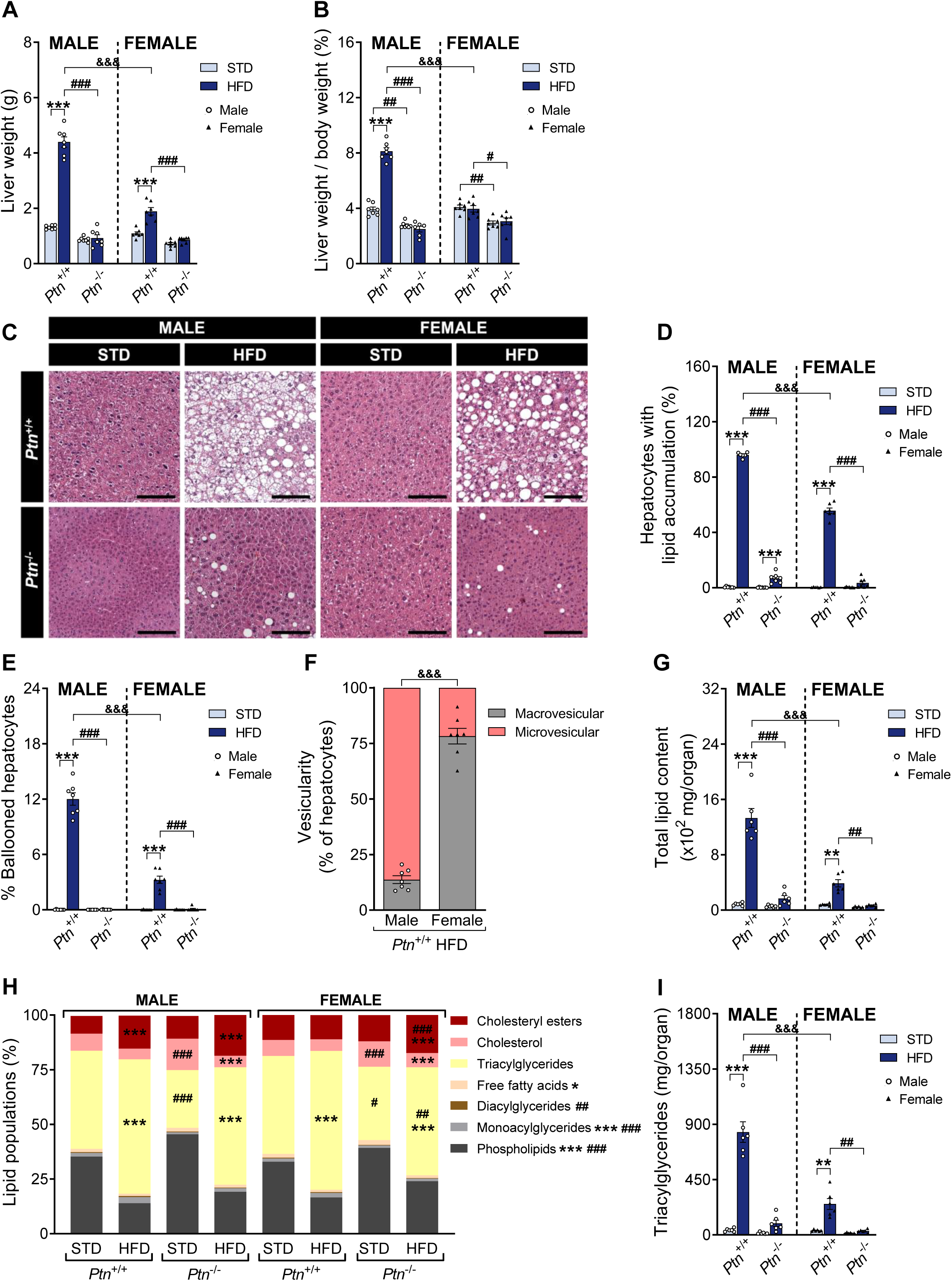
High-fat diet feeding and *Ptn* deletion effects on liver weight and steatosis. Liver weight (A); relative liver weight (B); hematoxylin-eosin staining of formalin-fixed paraffin-embedded liver tissue sections (C); percentage of hepatocytes with lipid accumulation (D); percentage of ballooned hepatocytes (E); characterisation of vesicularity on groups with steatosis (F); total lipid content by organ (G); percentage of lipid populations (H), and triacylglycerides in liver (I) in male and female *Ptn*^+/+^ and *Ptn*^−/−^ mice fed with a STD or HFD for 6 months. Data are presented as mean ± SEM (n = 7 animals per group; circles correspond to males and triangles correspond to females). Differences between diets are represented with: \**p* < 0.05; \*\**p* < 0.01; \*\*\**p* < 0.001; between genotypes with: ^#^*p* < 0.05; ^##^*p* < 0.01; ^###^*p* < 0.001; and between sexes with: ^&&&^*p* < 0.001. Scale bar 200 µm.

Histological analysis based on H&E staining allowed the study of the degree of hepatic steatosis in the different experimental groups (**Figure 4C**). Three-way ANOVA revealed that genotype, diet, and sex significantly affected the percentage of hepatocytes with lipid accumulation, and a significant interaction between the three variables was found (**Figure 4D**). Kruskal-Wallis analysis indicated that genotype and diet, but not sex, significantly affected the percentage of ballooned hepatocytes (**Figure 4E**). These data show that only *Ptn*^+/+^ animals fed HFD showed a clear state of hepatic steatosis. Following this observation, the type of vesicularity in those experimental groups with hepatic steatosis was analysed (**Figure 4F**). We found that after feeding HFD, male *Ptn*^+/+^ animals had a higher percentage of microvesicularity compared to female *Ptn*^+/+^ mice, which had a higher percentage of macrovesicularity (**Figure 4F**) (statistics in **Supplementary Table 1D**). Taken together, these results indicate that HFD promotes hepatic steatosis in *Ptn*^+/+^ animals, with male mice more affected than female mice. Surprisingly, we found that *Ptn*^−/−^ animals were protected from developing HFD-induced hepatic steatosis, the effect being more pronounced in females than in males.

### Effects of high-fat diet and *Ptn* deletion on different lipid species in the liver

Following the observed results, we studied the hepatic lipid content. Three-way ANOVA revealed that genotype, diet, and sex significantly affected the lipid content, although we did not detect significant interactions between variables (**Supplementary figure 3A**). In addition, Three-way ANOVA revealed that genotype, diet, and sex significantly affected the total lipid content in the whole organ, and a significant interaction between the three variables was found (**Figure 4G**). These results suggest that HFD leads to an increase in hepatic lipids in *Ptn*^+/+^ animals, which is more pronounced in males than in females. However, *Ptn*^−/−^ animals are protected from this HFD-induced hepatic lipid accumulation, regardless of the sex of the animal. We then tested whether the different types of lipids change in the same way (**Figure 4H**). Three-way ANOVA showed that genotype significantly affected the phospholipids, monoacylglycerides (MAG), diacylglycerides (DAG), cholesterol, TAG, and cholesteryl esters populations; and diet significantly affected the phospholipids, MAG, cholesterol, free fatty acids (FFA), TAG, and cholesteryl esters fractions. However, sex did not affect any lipid type population. Interaction between the three variables was found in the case of TAG and the cholesteryl esters population. To further explore the effect of genotype and diet, in cases where sex did not affect or interact with the other variables, we employed two-way ANOVA excluding the sex variable. Again, genotype significantly affected the phospholipids, MAG, DAG, and cholesterol population; and diet significantly affected the amount of phospholipids, MAG, cholesterol, and FFA. In addition, the interaction between the two variables was found in the case of cholesterol. In summary, HFD intake increased the percentage of cholesteryl esters, TAG and MAG, but decreased the percentage of phospholipids and FFA in *Ptn*^+/+^ animals. Moreover, the deletion of *Ptn* results in a decrease in the percentage of all acylglycerides in favour of an increase in the cholesterol and phospholipids. Finally, HFD partially restore the amount of the different lipid populations in *Ptn*^−/−^ animals to the values similar to those found in *Ptn*^+/+^ mice.

Finally, we studied each of the liver lipid species individually in order to determine the total lipid content in the animals studied (**Supplementary figure 3**). Three-way ANOVA showed that genotype, diet, and sex significantly affected the liver content of all lipid populations studied and a significant interaction between variables was found in all cases (statistics in **Supplementary Table 1D and K**). Briefly, HFD increased the liver content of all lipid populations in *Ptn*^+/+^ animals, which was more pronounced in male than in female mice. However, *Ptn* deletion protected against these HFD-induced alterations.

### *Ptn* deletion alters proteins involved in hepatic lipid metabolism, protecting against HFD-induced metabolic disruptions

The amount and phosphorylation of key proteins involved in lipid synthesis were studied to understand the molecular mechanism by which the observed hepatic lipid alterations occurred (**Figure 5**). Three-way ANOVA revealed that only diet significantly affected the AQP9 protein. In addition, no interaction between the three variables was found. To further explore the effect of genotype and diet, we employed a two-way ANOVA, excluding the sex variable. Again, we found a significant effect of the diet. However, a significant interaction between genotype and diet was found. In summary, *Ptn*^−/−^ animals were protected against the HFD-induced decrease in AQP9 observed in *Ptn*^+/+^ animals (**Figure 5A**).

**Figure 5.**
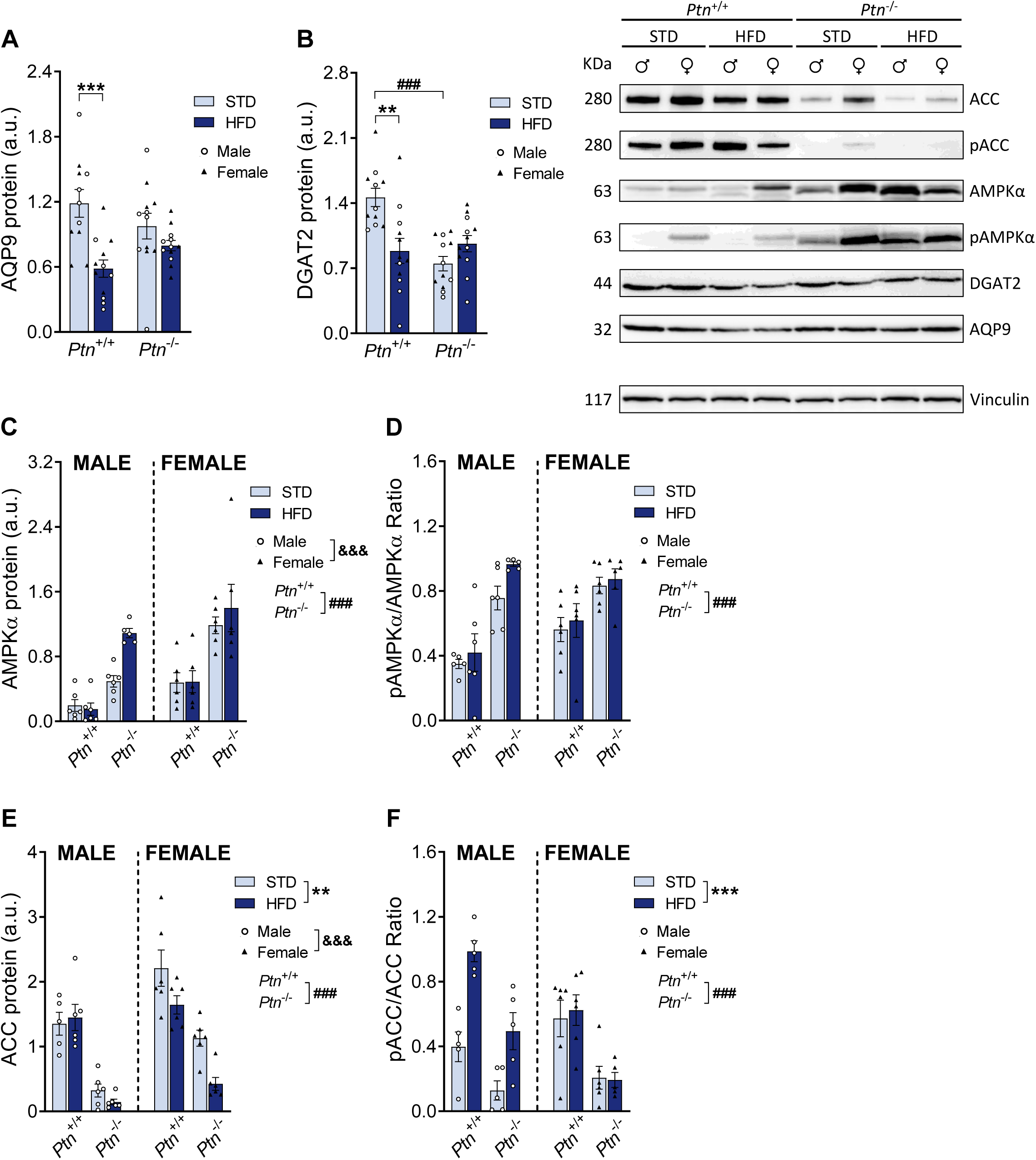
High-fat diet and *Ptn* deletion effects on the proteins involved in lipid synthesis in the liver. Aquaporin 9 (AQP9) (A); Diacylglycerol O-acyltransferase 2 (DGAT2) (B); AMP-activated protein kinase α (AMPKα) (C); phospho-AMPKα (pAMPKα)/AMPKα ratio (D); Acetyl-CoA carboxylase (ACC) hepatic protein level (E) and phospho-ACC (pACC)/ACC ratio (F) in male and female *Ptn*^+/+^ and *Ptn*^−/−^ mice fed with a STD or HFD for 6 months. Data are presented as mean ± SEM (n = 6 animals per group; circles correspond to males and triangles correspond to females). Differences between diets are represented with: \*\**p* < 0.01; \*\*\**p* < 0.001; between genotypes with: ^###^*p* < 0.001; and between sexes with: ^&&&^*p* < 0.001.

Next, a three-way ANOVA revealed that genotype significantly affected DGAT2 liver protein and a tendency was found for diet. In addition, no interaction between the three variables was found. To further explore the effect of genotype and diet, we employed two-way ANOVA excluding the sex variable. Again, we found a significant effect of the genotype and a tendency of the diet. A significant interaction between genotype and diet was found. *Post-hoc* analyses concluded that HFD is associated with a decrease of DGAT2 in *Ptn*^+/+^ mice. In addition, *Ptn*^−/−^ animals showed a decrease in DGAT2 that was not affected by HFD (**Figure 5B**).

Finally, we studied the AMPKα and ACC proteins and their phosphorylated forms and calculated the pACC/ACC and pAMPKα/AMPKα ratios (**Figures 5C-F**). For AMPKα, three-way ANOVA revealed that genotype and sex, but not diet, significantly affected the AMPKα total protein, but no interaction between the three variables was found (**Figure 5C**). Moreover, three-way ANOVA revealed that only genotype significantly affected the pAMPKα/AMPKα ratio. In addition, no interaction between the three variables was found (**Figure 5D**). For ACC, three-way ANOVA revealed that genotype, diet and sex significantly affected the ACC total protein, but no interaction between the three variables was found (**Figure 5E**). Moreover, three-way ANOVA revealed that genotype and diet, but not sex, significantly affected the pACC/ACC ratio. In addition, no interaction between the three variables was found (**Figure 5F**) (statistics in **Supplementary Table 1E**). We observed that females had higher AMPKα and ACC protein levels than male animals. In addition, HFD decreased ACC protein in female mice, but not in male mice. Finally, we found that, in the liver of *Ptn*^−/−^ mice, there was a drastic increase in AMPKα alongside a marked decrease in ACC. When studying AMPKα and ACC phosphorylation, we found that HFD increases the phosphorylation of ACC only in male mice. Again, we found that *Ptn*^−/−^ animals show a significant increase in pAMPKα/AMPKα ratio in contrast to a significant decrease in pACC/ACC ratio when compared to *Ptn*^+/+^ mice.

### *Ptn* deletion decreases collagen in the tunica adventitia of blood vessels and protects against the development of HFD-induced liver fibrosis

Next, to determine the degree of liver fibrosis, we studied total, type I and type III collagen of the tunica adventitia of blood vessels using picrosirius red collagen staining (**Figure 6A**). Three-way ANOVA indicated that genotype, diet and sex significantly affected the total, type I, and type III collagen (**Figure 6B-D**). Furthermore, a significant interaction between the three variables was found in the three cases. In addition, we also studied the proportions of type I and type III collagen (**Figure 6E**). Three-way ANOVA indicated that genotype and diet, but not sex, significantly affected the type I and type III collagen ratio. Moreover, a significant interaction between three variables was found only for type III collagen ratio (statistics in **Supplementary Table 1F**). Our data revealed that HFD favoured the development of liver fibrosis, increasing the total, type I, and type III collagen, and their relative proportions, only in male *Ptn*^+/+^ animals. Strikingly, the deletion of *Ptn* not only protected against this effect of HFD, but this animal model showed a decrease in collagen in the tunica adventitia of blood vessels, and the proportions were altered after HFD feeding.

**Figure 6.**
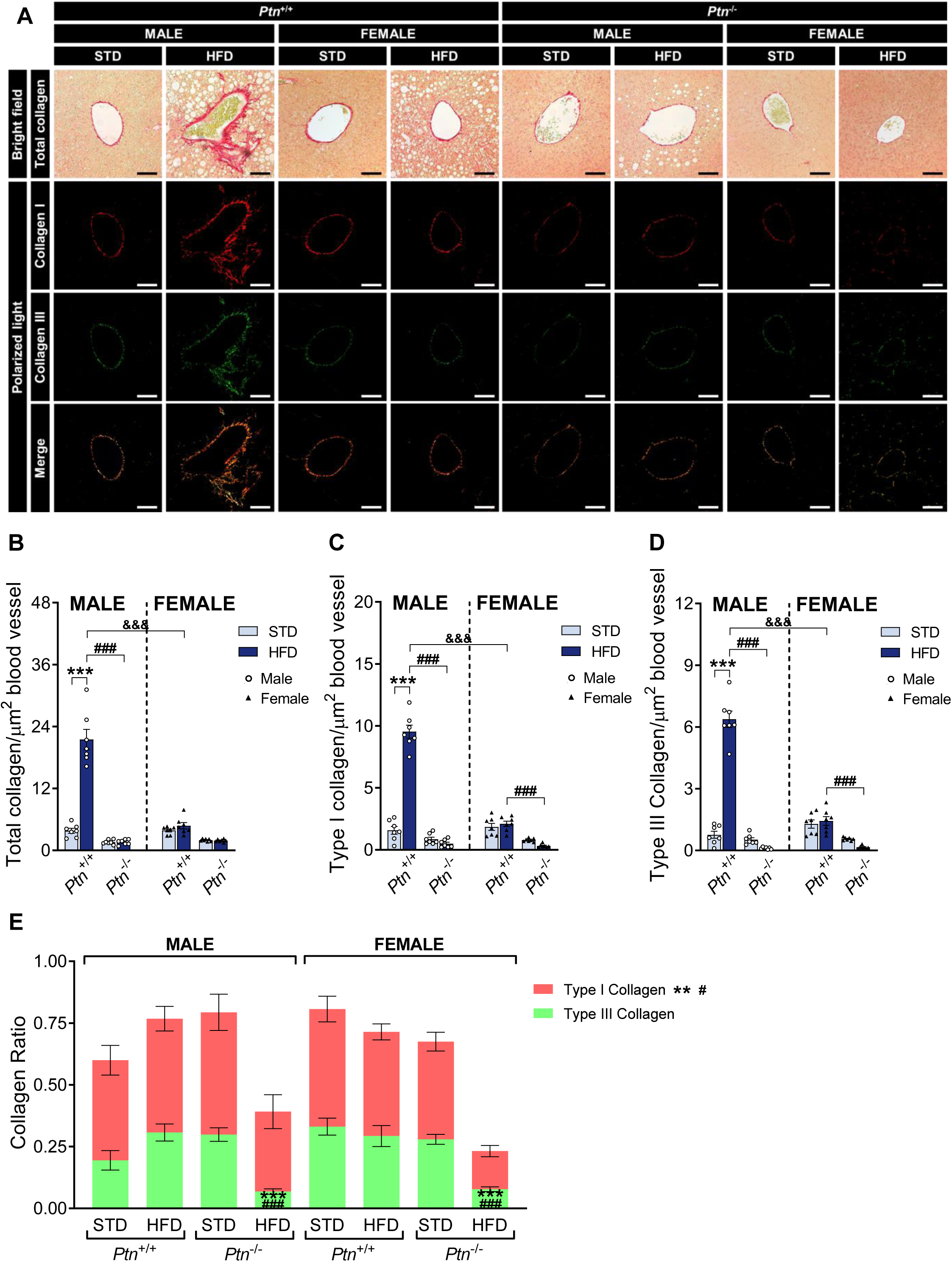
High-fat diet feeding and *Ptn* deletion effects on liver fibrosis. Picrosirius red staining of formalin-fixed paraffin-embedded liver tissue sections (A); marked area per µm^2^ of blood vessel of total (B), type I (C) and type III collagen (D); type I and type III collagen ratio versus total collagen (E) in male and female *Ptn*^+/+^ and *Ptn*^−/−^ mice fed with a STD or HFD for 6 months. Data are presented as mean ± SEM (n= 7 animals per group). Differences between diets are represented with: \**p* < 0.05; \*\*\**p* < 0.001; between genotypes with: ^##^*p* < 0.01; ^###^*p* < 0.001; and between sexes with: ^&^*p* < 0.05; ^&&&^*p* < 0.001. Scale bar 200 µm

### PTN induces triacylglyceride deposition in the human Huh7 cells by increasing AKT phosphorylation and reducing ACC phosphorylation

To understand the results observed *in vivo*, the effect of PTN on the human Huh7 hepatocyte cell line was studied. First, we examined the lipid accumulation in Huh7 cells by OR staining and TAG measurement (**Figure 7A**). Two-way ANOVA indicated that both insulin and PTN significantly increased both the lipid marked with OR and the TAG accumulation (**Figures 7B and C**). In addition, only a significant interaction between variables was found for the accumulated lipids marked with OR. Our results showed that PTN increased lipid accumulation and TAG synthesis at the same level as insulin. In addition, it also showed that its effect is greater when both signals are combined.

**Figure 7.**
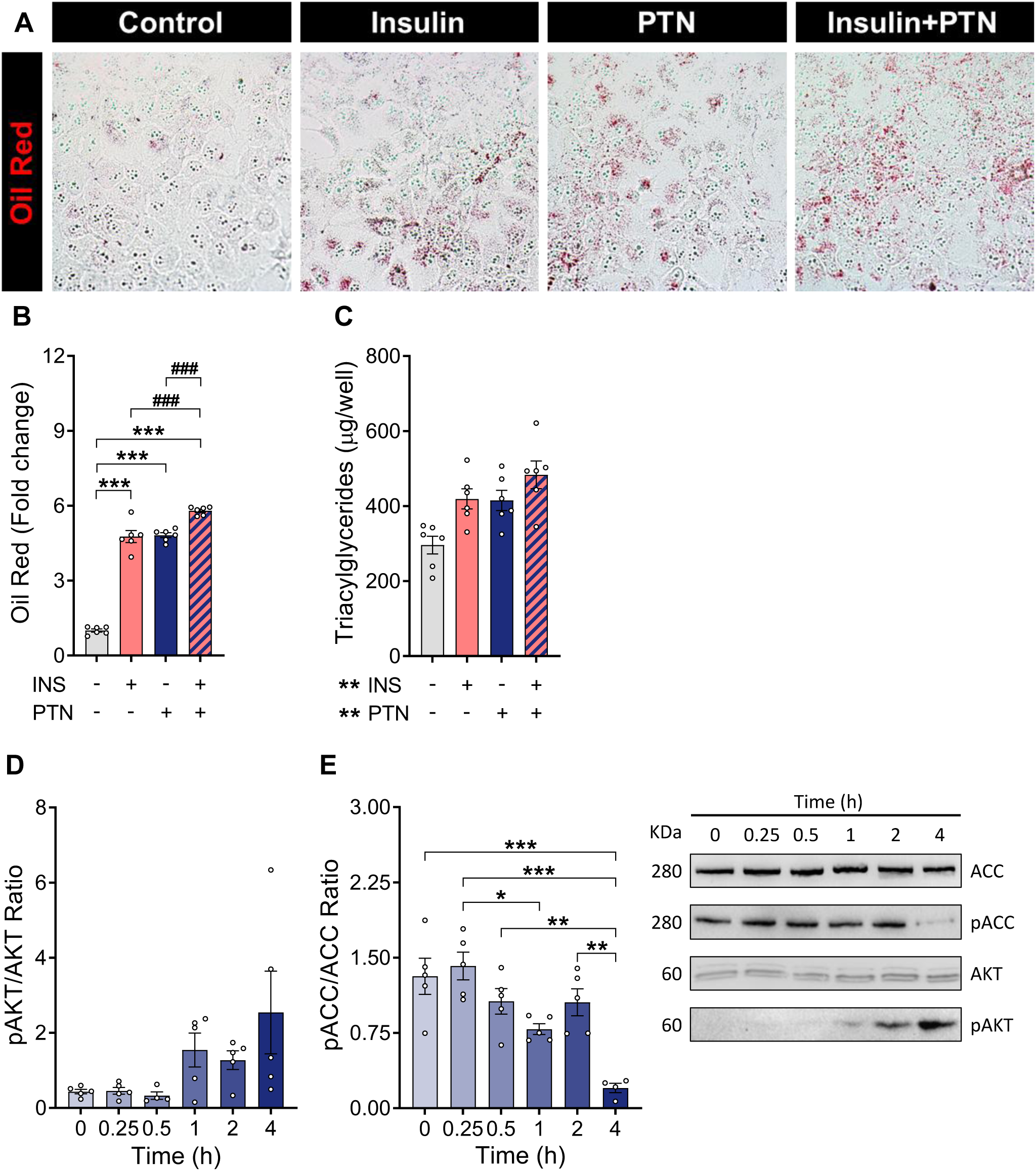
PTN effects on lipid accumulation and enzymes involved in lipid synthesis in Huh7 cell line. Oil Red staining of Huh7 cell line (A); fold change of accumulated lipids marker with Oil Red stain (B) and triacylglycerides per well (C) in Huh7 cell line treated with insulin and/or PTN for 48 hours. The treatment effects are represented with: \*\**p* < 0.01; \*\*\**p* < 0.001. Differences between treatments are represented with: ^###^*p* < 0.001. Phospho-protein kinase B (pAKT)/AKT ratio (D) and phospho-acetyl-CoA carboxylase (pACC)/ACC ratio (E) in Huh7 cell line treated with PTN and measured at different time points. Data are presented as mean ± SEM (n = 5-6 per group). Differences between time points are represented with: \**p* < 0.05; \*\**p* < 0.01; \*\*\**p* < 0.001.

We then analysed the effect of PTN on essential enzymes of lipid synthesis. One-way ANOVA revealed that PTN significantly increased the pAKT/AKT ratio while decreasing the pACC/ACC ratio (**Figures 7D and E**) (statistics in **Supplementary Table 1G**).

Finally, to elucidate the signalling pathways involved in these alterations, we analysed the mRNA of the known PTN receptors (**Figure 8**). Two-way ANOVA revealed that insulin significantly increased *ITGAV*, *LRP1*, *NRP1* and *SDC1* expression levels. However, no significant effects of the PTN treatment or interaction between the variables were observed (statistics in **Supplementary Table 1H**). Moreover, *ALK* and *PTPRZ1* mRNA levels were below the detection limits (data not shown).

**Figure 8.**
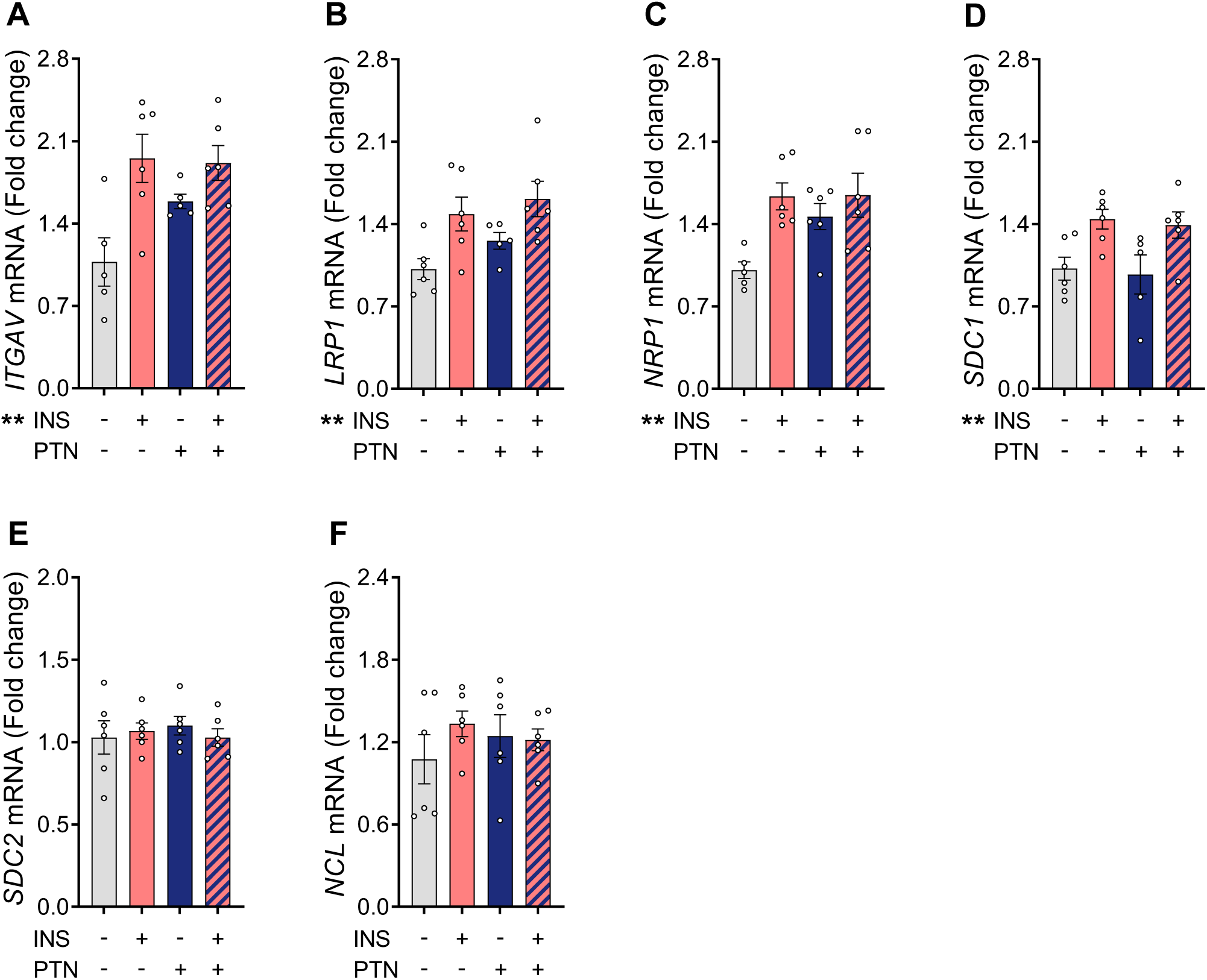
mRNA of PTN receptors in the Huh7 cell line. Integrin subunit alpha V (*ITGAV*) mRNA (A); LDL receptor related protein 1 (*LRP1*) mRNA (B); Neuropilin 1 (*NRP1*) mRNA (C); Syndecan 1 (*SDC1*) mRNA (D); Syndecan 2 (*SDC2*) mRNA (E) and Nucleolin (*NCL*) mRNA (F) in the Huh7 cell line treated with insulin and/or PTN for 48 hours. Anaplastic lymphoma kinase (*ALK*) mRNA and protein tyrosine phosphatase receptor type Z1 (*PTPRZ1*) mRNA levels were below the detection limits. Data are presented as mean ± SEM (n = 6 per group). The treatment effects are represented with: \*\**p* < 0.01.

## DISCUSSION

Obesity and its comorbidities, such as MetS and MASLD, are a global health problem of growing concern [1–3, 12, 27]. Among factors involved in these pathologies, chronic inflammation and metabolic imbalance are central elements that accelerate liver damage and progression to severe conditions such as MASH [9–11]. In this context, PTN, a cytokine that has been previously studied for its role in tissue regeneration and energy metabolism, is emerging as a potential regulator of liver homeostasis [21]. However, its specific role in liver function and obesity-related metabolic alterations has not been completely investigated.

As expected, HFD induced the development of obesity in *Ptn*^+/+^ animals, whereas *Ptn*^−/−^ animals did not gain weight even after 6 months on a 60% HFD. This confirms previous results in female mice with a less severe dietary treatment [5, 21]. Furthermore, our study demonstrates that the impact of *Ptn* deletion in mice fed a HFD for 6 months is consistent in both sexes. As there were no differences in the food intake between the groups, we can assess that *Ptn* deletion protects against HFD-induced body weight gain.

The expression in the liver of *Ptn* and *Mdk* has not been extensively studied, which renders findings of particular interest. We demonstrate that both *Ptn* and *Mdk* are expressed in the liver, with females showing higher *Mdk* expression than males. Importantly, HFD consumption alters the expression of both genes in a sexually dimorphic manner. This differential expression may play a significant role in the hepatic response to diet.

Obesity is closely associated with liver damage and impaired metabolic function, including gluconeogenesis. Our results suggest that obesity is associated with hyperactivation of gluconeogenesis, which is consistent with hepatic insulin resistance [28, 29]. In addition, we observed a sexual dimorphism, with females showing a greater capacity to normalise glycemia over time compared to males, who exhibited a continuous elevation of circulating glucose. This suggests an estrogen-mediated protective effect that may attenuate the metabolic dysfunction associated with insulin resistance [30]. By contrast, *Ptn*^−/−^ mice had an opposite metabolic profile, characterised by low basal glucose levels and reduced gluconeogenesis, suggesting that PTN acts as a positive regulator of hepatic gluconeogenesis. The hyperinsulinemia of *Ptn*^−/−^ mice observed in previous studies [21] may explain these effects, as elevated insulin levels can inhibit hepatic glucose production in the absence of insulin resistance [29, 31, 32]. In this context, *Ptn* deletion may enhance the ability of insulin to suppress gluconeogenesis, resulting in a better metabolic profile. The molecular events involved in the interaction between PTN and insulin in the modulation of hepatic gluconeogenesis could be mediated by AKT phosphorylation, as it will be discussed below. This alteration in liver metabolism impacts not only glucose homeostasis but also triggers a state of systemic inflammation, contributing to a vicious cycle of damage and dysfunction. Therefore, our results point to hepatic alterations and metainflammation with changes in the synthesis of plasma proteins in the liver. In fact, *Ptn*^+/+^ animals fed a HFD showed an increase in total α-globulin fractions and a decrease in albumin, indicating an inflammatory condition. In line with previous observations, this HFD-induced inflammatory state is less pronounced in female *Ptn*^+/+^ mice, indicating a delay in liver damage. Liver damage in *Ptn*^+/+^ animals may induce a negative feedback loop, where metabolic damage in the liver perpetuates a state of inflammation and alterations in other organs. This is relevant given the fundamental role of the liver in maintaining whole-body homeostasis, as damage to this organ can have detrimental consequences. However, *Ptn*^−/−^ animals fed a HFD did not show these alterations, suggesting a protective role of *Ptn* deletion. Furthermore, the analysis of circulating proinflammatory markers, such as TNF and CCL2, confirms the development of HFD-induced systemic inflammation which was absent in *Ptn*^−/−^ animals along with a reduced basal inflammatory state. These results align with previous studies that found modulation of systemic TNFα levels by endogenous PTN in both basal and LPS-induced inflammation [33]. Therefore, the present data suggest that physiological endogenous levels of PTN are required to trigger the systemic inflammatory response.

In addition, the electrophoretic analysis of circulating proteins revealed high presence of pre-albumin (also transthyretin), a transport protein for thyroid hormones [34], in *Ptn*^−/−^ mice. This result suggests a faster and more efficient transport of thyroid hormones to their target tissues and explains the decrease in circulating thyroxine in this animal model due to higher metabolism and uptake by the main target organs, as we have previously found [5, 20].

One of the most prevalent liver diseases associated with obesity is MASLD, which is characterised by excessive accumulation of fat in the liver, leading to lipotoxic damage. In this context, we observed that HFD led to a disproportionate increase in liver weight in male *Ptn*^+/+^ animals. This alteration is likely due to hepatic lipid accumulation in both sexes and an associated hepatic inflammation in the case of males. We found that *Ptn*^−/−^ animals had reduced liver size, possibly related to the mitogenic role of PTN, which did not change with HFD. Histological studies confirmed that HFD led to hepatic steatosis in *Ptn*^+/+^ animals, more pronounced in males, as shown by the increased lipid accumulation and ballooned hepatocytes, a marker of hepatocyte degeneration. In addition, males and females showed a differential distribution with predominantly microvesicular lipid accumulation in males and macrovesicular in females, supporting a sexual dimorphism in response to HFD. Analysis of hepatic lipid content revealed that HFD increased total lipid types and content, again more markedly in male *Ptn*^+/+^ animals. Surprisingly, *Ptn*^−/−^ mice did not show signs of hepatic steatosis or lipid accumulation, despite the prolonged duration and the high-fat content of the diet. Furthermore, when studying the lipid species, we found that *Ptn* deletion leads to a decrease in the percentage of all acylglycerides, with TAG being the major lipid specie, in favour of an increase in the cholesterol and phospholipids. These effects can be explained by alterations in key enzymes of hepatic lipid synthesis. The high lipid content and highly elevated TAG ratio in the liver of *Ptn*^+/+^ animals may lead to a decrease in AQP9 and DGAT2 as a compensatory mechanism to normalise lipid ratios. In contrast, *Ptn*^−/−^ animals show a constitutive decrease in DGAT2, an essential protein for hepatic TAG synthesis. A decrease or inhibition of this protein is associated with a reduction in hepatic fat and protects against HFD-induced hepatic steatosis [35, 36]. In this line of evidence, we found a drastic reduction in the amount of ACC, the rate-limiting enzyme of lipogenesis, which would support the lower fatty acid synthesis in the liver of these animals. Furthermore, although HFD was associated with an increase in inhibitory phosphorylation of ACC, probably as a mechanism to compensate for the excess hepatic lipids in males, this is not observed in females, regardless of the diet. In addition, we found that *Ptn*^−/−^ animals exhibited a significant increase in AMPKα protein levels and activation, as indicated by a higher pAMPKα/AMPKα ratio, in contrast to a marked decrease in total ACC levels mentioned above, and reduced pACC/ACC ratio. AMPKα acts as an energy sensor and a regulator of catabolic processes; when activated, it inhibits ACC, thereby reducing lipogenesis and promoting fatty acid oxidation [37]. The increased AMPKα activity in *Ptn*^−/−^ mice is consistent with their resistance to hepatic steatosis and suggests that PTN may negatively regulate AMPKα, either directly or indirectly, thereby promoting lipogenesis. The lower pACC/ACC ratio in *Ptn*^−/−^ mice could reflect an apparent disengagement of the AMPKα-ACC axis. However, this is likely due to the drastically reduced levels of total ACC protein in this model, which may mask the degree of phosphorylation and inhibition, and suggests a metabolic reprogramming that downregulates lipogenesis at both transcriptional and post-translational levels. This differential regulation of AMPKα and ACC, and the constitutive downregulation of DGAT2 in the *Ptn*^−/−^ animals, explains the protection against lipid accumulation and hepatic steatosis found in these animals. Furthermore, it also points to a sexual dimorphism in adaptive metabolic strategies, with females being more efficient in limiting lipid accumulation under HFD.

Interestingly, we found that *Ptn* was overexpressed only in the liver of male *Ptn*^+/+^ mice under HFD. This sex-specific increase is consistent with the higher degree of steatosis and inflammation observed in male mice. These findings, together with the protection observed in *Ptn*^−/−^ animals, suggest that basal PTN levels are necessary for the damaging effects of the HFD. However, the lack of response in females may be indicative of their enhanced metabolic resilience, as has been previously suggested. In parallel, *Mdk*, a cytokine homologous to PTN and frequently co-regulated, exhibited a remarkably similar pattern to *Ptn.* Specifically, the overexpression of *Mdk* was observed only in the liver of *Ptn*^+/+^ males fed with HFD. In contrast, females showed higher baseline *Mdk* mRNA levels, which remained constant despite the intake of the HFD. This finding suggests a potential mechanism contributing to their reduced susceptibility to developing steatosis and inflammation. In turn, our *in vitro* results support this hypothesis, showing that PTN induces an increase in TAG levels and a lipid accumulation profile similar to the lipogenic effect of insulin. Moreover, we observed that the effects of PTN are significantly enhanced when combined with insulin, suggesting that both molecules share non-competitive signalling pathways. PTN is a ligand of the syndecan receptors [38, 39], which trigger phosphorylation and activation of AKT [40–42], promoting lipid synthesis in hepatocytes. In addition, we found that PTN reduces the phosphorylation of ACC, favouring the active form of the enzyme due to the inhibition of AMPKα. This mechanism integrates well with that observed in *Ptn*^−/−^ animals, where the absence of PTN disrupts these pathways protecting against hepatic steatosis.

All of the above described alterations lead to progressive liver damage, which in turn triggers a state of liver fibrosis. This condition, in which the liver is already at an advanced stage of the disease, is known as MASH [9, 12]. This process involves not only lipid accumulation but also significant changes in the composition and organisation of the extracellular matrix (ECM), particularly in the tunica adventitia of the hepatic blood vessels [43, 44]. Our results showed that in male *Ptn*^+/+^ mice, HFD induced excessive remodelling of the ECM characteristic of liver fibrosis. This increase in total collagen was accompanied by higher proportions of type I and III collagens, which are closely associated with the structural deterioration characteristic of this advanced disease stage [44]. In contrast, *Ptn* deletion protected against the development of HFD-induced liver fibrosis and reduced total collagen levels in the tunica adventitia of blood vessels. *Ptn*^−/−^ animals showed a significant decrease in the ratios of collagen types I and III after HFD treatment. This suggests that the absence of PTN interferes with signalling pathways responsible for ECM synthesis and remodelling, preventing excessive collagen accumulation and, consequently, mitigating the progression to liver fibrosis. Again, the differences observed between males and females underline the influence of sex on the liver-damaging effects of HFD. In this context, the relationship between PTN and ACC seems to play a key role. Other studies suggest that inhibition of ACC significantly reduces fibrosis in various MASH models [45]. In this line of evidence, PTN may increase collagen production and promote HFD-induced liver fibrosis through ACC activation. This is because ACC activity culminates in the development of lipotoxicity, which promotes inflammatory signals and the activation of hepatic stellate cells (HSCs), leading to fibrosis [46]. However, in the absence of PTN, total ACC levels are reduced and thus its activity. This, together with the decrease in inflammatory signals, may reduce collagen production through HSCs activation. This is consistent with the protection observed in this animal model and the lower levels of collagen under physiological conditions.

Overall, this study is of significant importance in the context of the increasing global prevalence of obesity and MetS, key factors in the development of chronic diseases such as MASLD and MASH. It identifies PTN as a key regulator in metabolic, inflammatory and liver remodelling processes linked to obesity and MetS. *Ptn* deletion protects against HFD-induced weight gain, systemic inflammation, hepatic lipid accumulation and the development of steatosis and liver fibrosis. Furthermore, our study also points to a sexual dimorphism in adaptive metabolic strategies, with females demonstrating a greater degree of protection against the liver-damaging effects of the HFD. These findings contribute to our understanding of the underlying molecular mechanisms and point to PTN as a promising therapeutic target to address the global problem of these metabolic diseases.

## DECLARATIONS

### Availability of data and materials

The datasets used and/or analysed during the current study are available from the corresponding author upon reasonable request.

### Consent for publication

Not applicable.

## Authors’ contributions

**HC-R**: Methodology, Investigation, Data Curation, Formal analysis, Data Interpretation, Visualization, Writing - Original Draft, Writing - Review and Editing. **AZ**; **MIS-C**; **FR; AO-R** and **JP-D**: Methodology, Investigation, Data Curation. **ML**; **JP-S**; **MGS-A**; **JS**; **A.M-V** and **EG**: Methodology, Investigation. **GH**: Conceptualization, Methodology, Funding acquisition, Data Interpretation, Visualization, Validation, Supervision, Project administration, Writing - Review and Editing. **MPR-A**: Conceptualization, Methodology, Funding acquisition, Data Interpretation, Visualization, Validation, Supervision, Project administration, Writing - Original Draft, Writing - Review and Editing. All authors have read and agreed to the published version of the manuscript.

## Acknowledgements

HC-R was supported by a fellowship from Fundación Universitaria San Pablo CEU-Santander. MIS-C was supported by a fellowship from Comunidad de Madrid. We thank our colleagues in the animal facility at the Universidad San Pablo CEU.

---

**Supplementary Table 1A.** Statistical analysis from Figure 1.

| Test | Genotype | Diet | Interaction |  |
| --- | --- | --- | --- | --- |
| Two-way repeated measures ANOVA (A) | $F_{1, 24} = 75.961; p < 0.001$ | $F_{1, 24} = 193.314; p < 0.001$ | $F_{1, 24} = 58.525; p < 0.001$ | |
| Two-way repeated measures ANOVA (B) | $F_{1, 24} = 37.078; p < 0.001$ | $F_{1, 24} = 105.375; p < 0.001$ | $F_{1, 24} = 40.741; p < 0.001$ | |
|  | Genotype | Diet | Sex | Interaction |
| Three-way ANOVA (C) | $F_{1, 55} = 108.697; p < 0.001$ | $F_{1, 55} = 129.295; p < 0.001$ | $F_{1, 55} = 32.333; p < 0.001$ | n.s. |
| Three-way ANOVA (E) | $F_{1, 53} = 133.666; p < 0.001$ | $F_{1, 53} = 78.995; p < 0.001$ | $F_{1, 53} = 13.677; p = 0.001$ | n.s. |
|  | Genotype | Diet | Interaction |  |
| Two-way repeated measures ANOVA (F) | $F_{1, 24} = 107.414; p < 0.001$ | $F_{1, 24} = 54.787; p < 0.001$ | $F_{1, 24} = 25.200; p < 0.001$ | |
| Two-way repeated measures ANOVA (G) | $F_{1, 23} = 104.806; p < 0.001$ | $F_{1, 23} = 24.080; p < 0.001$ | $F_{1, 23} = 21.410; p < 0.001$ | |
|  | Genotype | Diet | Sex | Interaction |
| Three-way ANOVA (H) | $F_{1, 53} = 236.007; p < 0.001$ | $F_{1, 53} = 76.982; p < 0.001$ | $F_{1, 53} = 12.083; p = 0.001$ | n.s. |

**Supplementary Table 1B.**
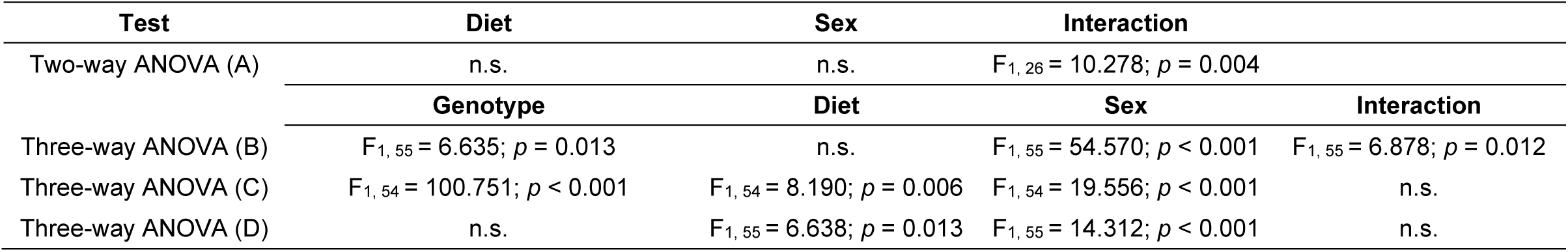
Statistical analysis from Figure 2.

| Test | Diet | Sex | Interaction |  |
| --- | --- | --- | --- | --- |
| Two-way ANOVA (A) | n.s. | n.s. | $F_{1,26} = 10.278; p = 0.004$ | |
|  | Genotype | Diet | Sex | Interaction |
| Three-way ANOVA (B) | $F_{1,55} = 6.635; p = 0.013$ | n.s. | $F_{1,55} = 54.570; p < 0.001$ | $F_{1,55} = 6.878; p = 0.012$ |
| Three-way ANOVA (C) | $F_{1,54} = 100.751; p < 0.001$ | $F_{1,54} = 8.190; p = 0.006$ | $F_{1,54} = 19.556; p < 0.001$ | n.s. |
| Three-way ANOVA (D) | n.s. | $F_{1,55} = 6.638; p = 0.013$ | $F_{1,55} = 14.312; p < 0.001$ | n.s. |

**Supplementary Table 1C.** Statistical analysis from Figure 3.

| Test | Genotype | Diet | Sex | Interaction |
| --- | --- | --- | --- | --- |
| Three-way ANOVA (A) | n.s. | $F_{1,47} = 5.179; p = 0.028$ | n.s. | $F_{1,47} = 4.555; p = 0.039$ |
| Kruskal-Wallis; Pre-Albumin (B.1) | **Genotype** | **Diet** | **Sex** |  |
| | $H(1) = 18.161; p < 0.001$ | $H(1) = 11.585; p = 0.001$ | n.s. | |
|  | **Genotype** | **Diet** | **Sex** | **Interaction** |
| Three-way ANOVA; Albumin (B.2) | n.s. | $F_{1,46} = 21.097; p < 0.001$ | n.s. | n.s. |
| Three-way ANOVA; $\alpha_1$-Globulin (B.3) | $F_{1,47} = 5.259; p = 0.027$ | $F_{1,47} = 9.674; p = 0.003$ | $F_{1,47} = 13.226; p = 0.001$ | n.s. |
| Three-way ANOVA; $\alpha_2$-Globulin (B.4) | n.s. | $F_{1,46} = 23.657; p < 0.001$ | $F_{1,46} = 41.660; p < 0.001$ | $F_{1,46} = 4.972; p = 0.032$ |
| Three-way ANOVA; β-Globulin (B.5) | n.s. | n.s. | n.s. | n.s. |
| Three-way ANOVA; γ-Globulin (B.6) | n.s. | n.s. | $F_{1,46} = 8.305; p = 0.006$ | n.s. |
| Kruskal-Wallis (C) | **Genotype** | **Diet** | **Sex** |  |
| | $H(1) = 18.161; p < 0.001$ | $H(1) = 12.157; p < 0.001$ | n.s. | |
|  | **Genotype** | **Diet** | **Sex** | **Interaction** |
| Three-way ANOVA (D) | $F_{1,47} = 7.357; p = 0.01$ | $F_{1,47} = 15.061; p < 0.001$ | $F_{1,47} = 7.329; p = 0.01$ | n.s. |
| Three-way ANOVA (E) | $F_{1,55} = 9.920; p = 0.003$ | $F_{1,55} = 20.955; p < 0.001$ | n.s. | n.s. |
| Two-way ANOVA (E') | **Genotype** | **Diet** | **Interaction** |  |
| | $F_{1,55} = 10.417; p = 0.002$ | $F_{1,55} = 22.005; p < 0.001$ | $F_{1,55} = 17.983; p < 0.001$ | |
|  | **Genotype** | **Diet** | **Sex** | **Interaction** |
| Three-way ANOVA (F) | $F_{1,55} = 143.726; p < 0.001$ | $F_{1,55} = 23.365; p < 0.001$ | n.s. | n.s. |
| Two-way ANOVA (F') | **Genotype** | **Diet** | **Interaction** |  |
| | $F_{1,55} = 140.886; p < 0.001$ | $F_{1,55} = 22.903; p < 0.001$ | $F_{1,55} = 16.961; p < 0.001$ | |

**Supplementary Table 1D.** Statistical analysis from Figure 4.

| Test | Genotype | Diet | Sex | Interaction |
| --- | --- | --- | --- | --- |
| Three-way ANOVA (A) | $F_{1,55} = 376.119; p < 0.001$ | $F_{1,55} = 221.875; p < 0.001$ | $F_{1,55} = 115.149; p < 0.001$ | $F_{1,55} = 74.624; p < 0.001$ |
| Three-way ANOVA (B) | $F_{1,54} = 260.181; p < 0.001$ | $F_{1,54} = 36.032; p < 0.001$ | $F_{1,54} = 51.618; p < 0.001$ | $F_{1,54} = 75.264; p < 0.001$ |
| Three-way ANOVA (D) | $F_{1,53} = 2319.832; p < 0.001$ | $F_{1,53} = 2990.280; p < 0.001$ | $F_{1,53} = 219.541; p < 0.001$ | $F_{1,53} = 155.799; p < 0.001$ |
| Kruskal-Wallis (E) | **Genotype** | **Diet** | **Sex** |  |
| | $H(1) = 15.626; p < 0.001$ | $H(1) = 15.549; p < 0.001$ | n.s. | |
|  | **Sex** |  |  |  |
| T-Student (F) | $t_{13} = 5.029; p < 0.001$ | | | |
|  | **Genotype** | **Diet** | **Sex** | **Interaction** |
| Three-way ANOVA (G) | $F_{1,47} = 103.502; p < 0.001$ | $F_{1,47} = 123.361; p < 0.001$ | $F_{1,47} = 48.587; p < 0.001$ | $F_{1,47} = 30.877; p < 0.001$ |
| Three-way ANOVA; Cholesteryl esters (H.1) | $F_{1,46} = 32.502; p < 0.001$ | $F_{1,46} = 69.387; p < 0.001$ | n.s. | $F_{1,46} = 7.804; p = 0.008$ |
| Three-way ANOVA; Cholesterol (H.2) | $F_{1,47} = 22.165; p < 0.001$ | $F_{1,47} = 53.043; p < 0.001$ | n.s. | n.s. |
| Three-way ANOVA; Triacylglycerides (H.3) | $F_{1,45} = 59.750; p < 0.001$ | $F_{1,45} = 139.361; p < 0.001$ | n.s. | $F_{1,45} = 4.175; p = 0.048$ |
| Three-way ANOVA; Free fatty acids (H.4) | n.s. | $F_{1,45} = 4.972; p = 0.032$ | n.s. | n.s. |
| Three-way ANOVA; Diacylglycerides (H.5) | $F_{1,45} = 10.618; p = 0.002$ | n.s. | n.s. | n.s. |
| Three-way ANOVA; Monoacylglycerides (H.6) | $F_{1,46} = 25.626; p < 0.001$ | $F_{1,46} = 16.448; p < 0.001$ | n.s. | n.s. |
| Three-way ANOVA; Phospholipids (H.7) | $F_{1,47} = 22.988; p < 0.001$ | $F_{1,47} = 169.335; p < 0.001$ | n.s. | n.s. |
|  | **Genotype** | **Diet** | **Interaction** |  |
| Two-way ANOVA; Cholesterol (H.2') | $F_{1,47} = 21.499; p < 0.001$ | $F_{1,47} = 51.218; p < 0.001$ | $F_{1,47} = 12.279; p = 0.001$ | |
| Two-way ANOVA; Free fatty acids (H.4') | n.s. | $F_{1,45} = 5.680; p = 0.022$ | n.s. | |
| Two-way ANOVA; Diacylglycerides (H.5') | $F_{1,45} = 11.303; p = 0.002$ | n.s. | n.s. | |
| Two-way ANOVA; Monoacylglycerides (H.6') | $F_{1,46} = 24.066; p < 0.001$ | $F_{1,46} = 14.630; p < 0.001$ | n.s. | |
| Two-way ANOVA; Phospholipids (H.7') | $F_{1,47} = 21.050; p < 0.001$ | $F_{1,47} = 155.055; p < 0.001$ | n.s. | |
|  | **Genotype** | **Diet** | **Sex** | **Interaction** |
| Three-way ANOVA (I) | $F_{1,47} = 104.262; p < 0.001$ | $F_{1,47} = 126.893; p < 0.001$ | $F_{1,47} = 43.558; p < 0.001$ | $F_{1,47} = 27.712; p < 0.001$ |

**Supplementary Table 1E.** Statistical analysis from Figure 5.

| Test | Genotype | Diet | Sex | Interaction |
| --- | --- | --- | --- | --- |
| Three-way ANOVA (A) | n.s. | $F_{1,46} = 17.287; p < 0.001$ | n.s. | n.s. |
|  | Genotype | Diet | Interaction |  |
| Two-way ANOVA (A') | n.s. | $F_{1,46} = 16.037; p < 0.001$ | $F_{1,46} = 4.685; p = 0.036$ | |
|  | Genotype | Diet | Sex | Interaction |
| Three-way ANOVA (B) | $F_{1,46} = 9.623; p = 0.004$ | $F_{1,46} = 3.053; p = 0.088$ | n.s. | n.s. |
|  | Genotype | Diet | Interaction |  |
| Two-way ANOVA (B') | $F_{1,46} = 9.650; p = 0.003$ | $F_{1,46} = 3.107; p = 0.085$ | $F_{1,46} = 14.950; p < 0.001$ | |
|  | Genotype | Diet | Sex | Interaction |
| Three-way ANOVA (C) | $F_{1,46} = 52.619; p < 0.001$ | n.s. | $F_{1,46} = 17.014; p < 0.001$ | n.s. |
| Three-way ANOVA (D) | $F_{1,45} = 46.641; p < 0.001$ | n.s. | n.s. | n.s. |
| Three-way ANOVA (E) | $F_{1,46} = 103.876; p < 0.001$ | $F_{1,46} = 8.846; p = 0.005$ | $F_{1,46} = 22.149; p < 0.001$ | n.s. |
| Three-way ANOVA (F) | $F_{1,42} = 39.783; p < 0.001$ | $F_{1,46} = 8.846; p = 0.005$ | n.s. | n.s. |

**Supplementary Table 1F.** Statistical analysis from Figure 6.

| Test | Genotype | Diet | Sex | Interaction |
| --- | --- | --- | --- | --- |
| Three-way ANOVA (B) | $F_{1,55} = 159.488; p < 0.001$ | $F_{1,55} = 75.980; p < 0.001$ | $F_{1,55} = 56.668; p < 0.001$ | $F_{1,55} = 61.836; p < 0.001$ |
| Three-way ANOVA (C) | $F_{1,55} = 292.150; p < 0.001$ | $F_{1,55} = 99.870; p < 0.001$ | $F_{1,55} = 101.040; p < 0.001$ | $F_{1,55} = 106.376; p < 0.001$ |
| Three-way ANOVA (D) | $F_{1,55} = 256.394; p < 0.001$ | $F_{1,55} = 86.523; p < 0.001$ | $F_{1,55} = 66.864; p < 0.001$ | $F_{1,55} = 106.413; p < 0.001$ |
| Three-way ANOVA; Type I collagen ratio (E.1) | $F_{1,55} = 7.118; p = 0.01$ | $F_{1,55} = 7.732; p = 0.008$ | n.s. | n.s. |
| Three-way ANOVA; Type III collagen ratio (E.2) | $F_{1,55} = 22.709; p < 0.001$ | $F_{1,55} = 18.079; p < 0.001$ | n.s. | $F_{1,55} = 4.536; p = 0.038$ |

**Supplementary Table 1G.**
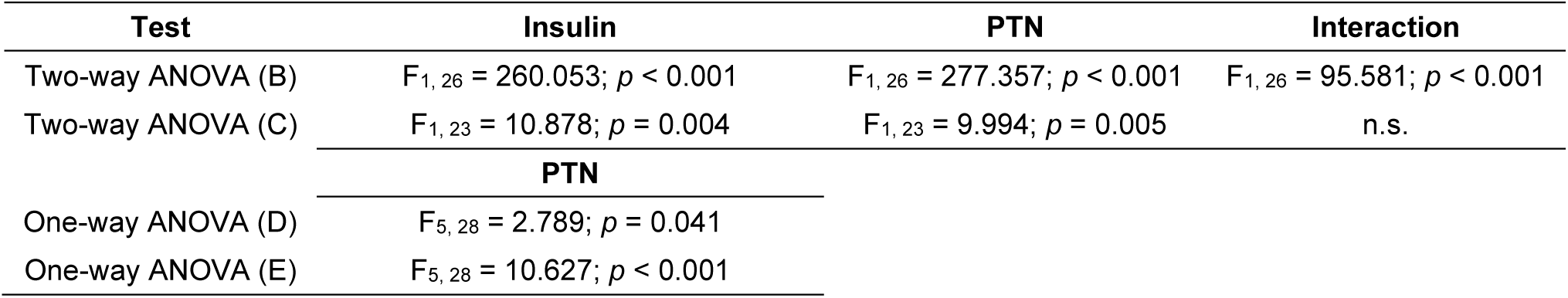
Statistical analysis from Figure 7.

**Supplementary Table 1H.** Statistical analysis from Figure 8.

| Test | Insulin | PTN | Interaction |
| --- | --- | --- | --- |
| Two-way ANOVA (A) | $F_{1,21} = 12.595; p < 0.001$ | n.s. | n.s. |
| Two-way ANOVA (B) | $F_{1,22} = 11.195; p = 0.003$ | n.s. | n.s. |
| Two-way ANOVA (C) | $F_{1,22} = 9.297; p = 0.007$ | n.s. | n.s. |
| Two-way ANOVA (D) | $F_{1,22} = 13.453; p = 0.002$ | n.s. | n.s. |
| Two-way ANOVA (E) | n.s. | n.s. | n.s. |
| Two-way ANOVA (F) | n.s. | n.s. | n.s. |

**Supplementary Table 1I.** Statistical analysis from Supplementary figure 1.

| Test | Genotype | Diet | Interaction |
| --- | --- | --- | --- |
| Two-way repeated measures ANOVA (A) | n.s. | $F_{1,21} = 14.218; p = 0.001$ | n.s. |
| Two-way repeated measures ANOVA (B) | n.s. | $F_{1,24} = 3.912; p = 0.06$ | n.s. |

**Supplementary Table 1J.** Statistical analysis from Supplementary figure 2.

| Test | Genotype | Diet | Sex | Interaction |
| --- | --- | --- | --- | --- |
| Three-way ANOVA (A) | n.s. | n.s. | n.s. | n.s. |
| Three-way ANOVA (B) | $F_{1,47} = 4.383; p = 0.043$ | $F_{1,47} = 16.813; p < 0.001$ | $F_{1,47} = 10.543; p = 0.002$ | $F_{1,47} = 9.058; p = 0.005$ |
| Three-way ANOVA (C) | $F_{1,47} = 7.138; p = 0.011$ | $F_{1,47} = 21.144; p < 0.001$ | $F_{1,47} = 12.104; p = 0.001$ | $F_{1,47} = 8.606; p = 0.006$ |
| Three-way ANOVA (D) | n.s. | n.s. | n.s. | n.s. |
| Three-way ANOVA (E) | n.s. | n.s. | $F_{1,46} = 7.524; p = 0.009$ | n.s. |

**Supplementary Table 1K.**
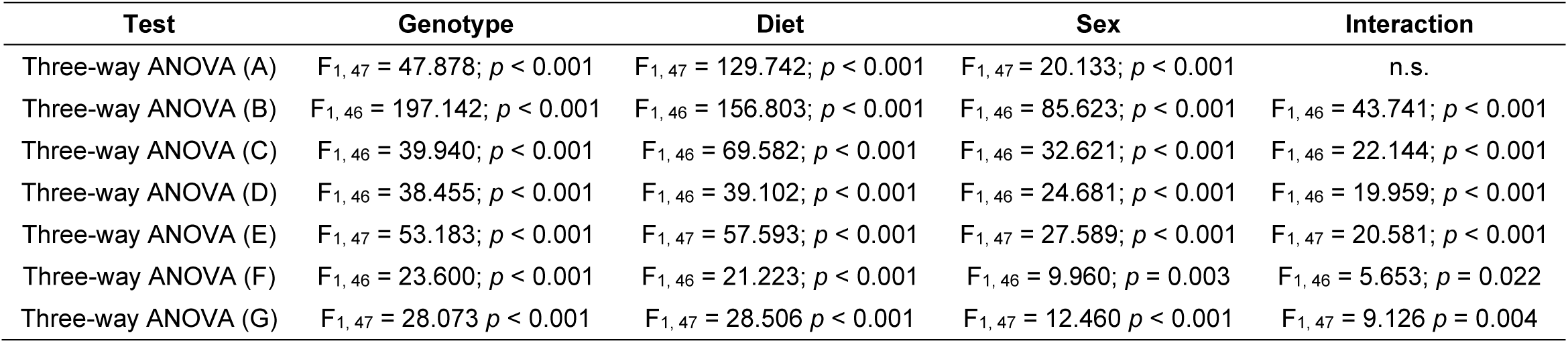
Statistical analysis from Supplementary figure 3.

**Supplementary figure 1.**
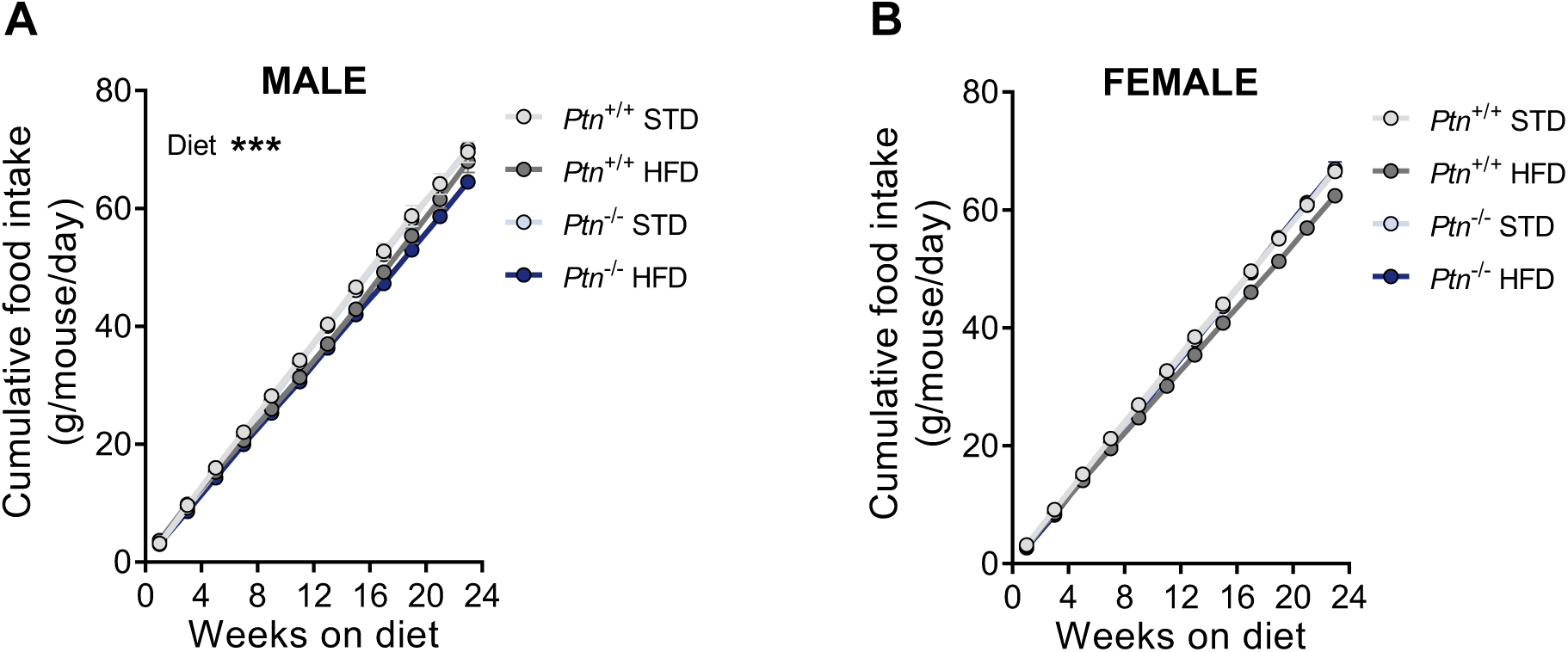
High-fat diet feeding and *Ptn* deletion effects on food intake. Cumulative food intake over time in male (A) and female (B) *Ptn*^+/+^ and *Ptn*^−/−^ mice fed with a STD or HFD for 6 months. Data are presented as mean ± SEM (n = 7 animals per group). Differences between diets are represented with: \*\*\**p* < 0.001.

**Supplementary figure 2.**
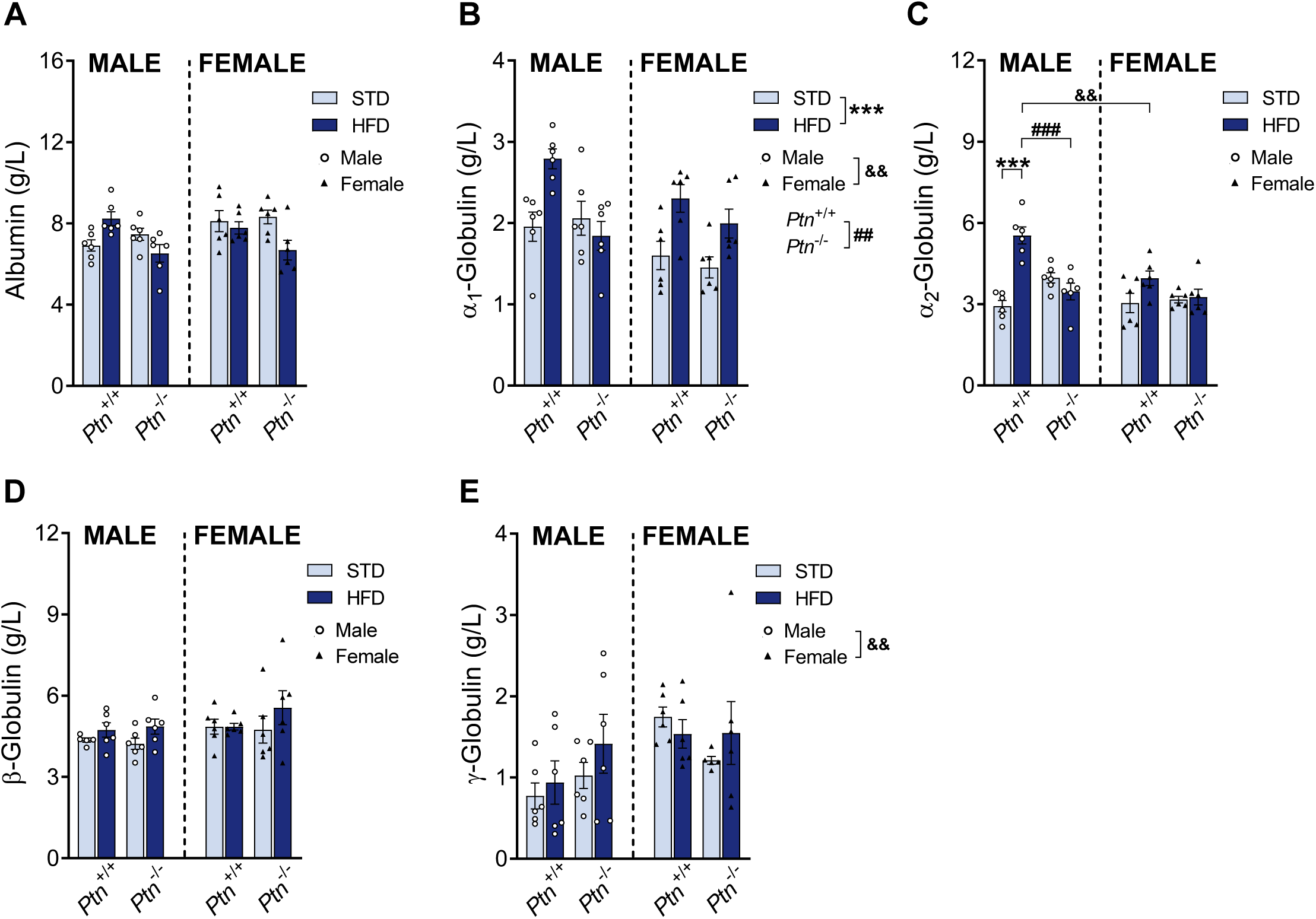
High-fat diet feeding and *Ptn* deletion effects on serum proteins. Albumin (A); α_1_-globulin fraction (B); α_2_-globulin fraction (C); β-globulin fraction (D), and γ-globulin fraction (E) in male and female *Ptn*^+/+^ and *Ptn*^−/−^ mice fed with a STD or HFD for 6 months. Data are presented as mean ± SEM (n= 6 animals per group). Differences between diets are represented with: \*\*\**p* < 0.001; between genotypes with: ^##^*p* < 0.01; ^###^*p* < 0.001; and between sexes with: ^&&^*p* < 0.01.

**Supplementary figure 3.**
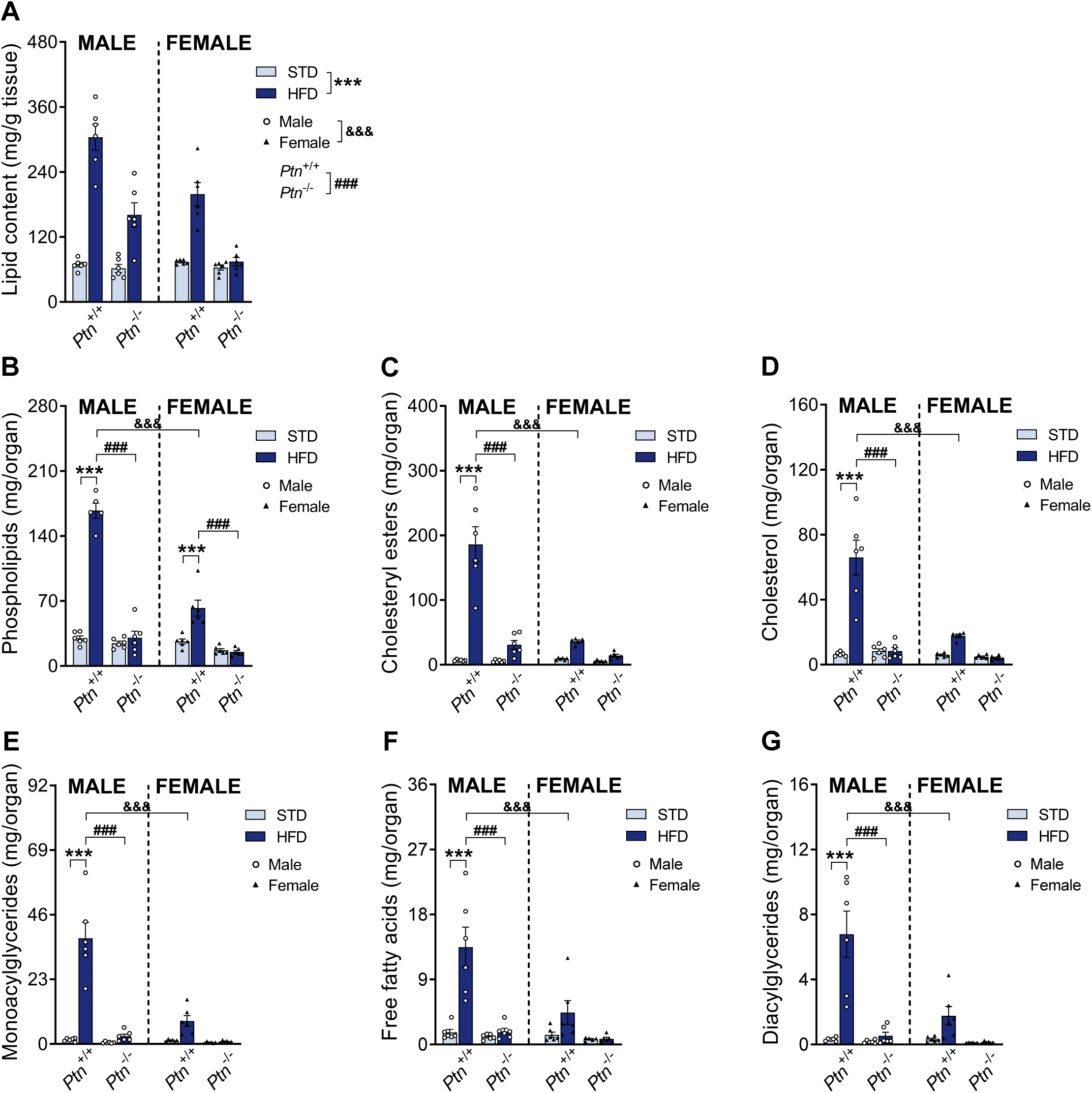
High-fat diet feeding and *Ptn* deletion effects on hepatic lipid content. Lipid content (A); phospholipids (B); cholesteryl esters (C); cholesterol (D); monoacylglycerides (E); free fatty acids (F), and diacylglycerides (G) content in the liver of male and female *Ptn*^+/+^ and *Ptn*^−/−^ mice fed with a STD or HFD for 6 months. Data are presented as mean ± SEM (n = 7 animals per group; circles correspond to males and triangles correspond to females). Differences between diets are represented with: \*\*\**p* < 0.001; between genotypes with: ^###^*p* < 0.001; and between sexes with: ^&&&^*p* < 0.001.

